# PCNA recruits cohesin loader Scc2/NIPBL to ensure sister chromatid cohesion

**DOI:** 10.1101/2022.09.16.508217

**Authors:** Ivan Psakhye, Ryotaro Kawasumi, Takuya Abe, Kouji Hirota, Dana Branzei

## Abstract

Sister chromatid cohesion is essential for faithful chromosome segregation and genome duplication during cell division. Failure to establish cohesion during S phase by the ring-shaped multiprotein complex cohesin leads to genomic instability^1-4^. Replisome-associated proteins are required to generate cohesion by two independent pathways^5^. One mediates conversion of cohesins bound to unreplicated DNA ahead of replication forks into cohesive entities behind them, while the second promotes cohesin de novo loading onto newly-replicated DNAs^6^. The latter process depends on the cohesin loader Scc2/NIPBL and the alternative PCNA loader CTF18-RFC. However, the precise mechanism of de novo cohesin loading during replication is unknown. Here we show that PCNA physically recruits yeast cohesin loader Scc2 via its C-terminal PCNA-interacting protein motif. Binding to PCNA is crucial, as *scc2-pip* mutant deficient in Scc2-PCNA interaction is defective in cohesion when combined with replisome mutants of the cohesin conversion pathway. Moreover, *scc2-pip* mutant becomes inviable without its partner Scc4/MAU2 that localizes cohesin loader to centromeres. Importantly, the role of NIPBL recruitment to PCNA for cohesion generation is conserved in vertebrate cells. Our results demonstrate that PCNA, the maestro of replication-linked functions, is also crucially involved in the cohesion establishment through de novo cohesin loading onto replicated DNA.

## Main

Sister chromatid cohesion (SCC) is mediated by cohesin, one of the three structural maintenance of chromosomes (SMC) complexes present in eukaryotic cells^1^. Similar to other SMC complexes, cohesin is composed of a pair of rod-shaped SMC proteins, Smc1 and Smc3, that heterodimerize via their hinge domains and are connected by a flexible kleisin subunit Scc1 at their ATPase head domains, thus forming heterotrimeric rings. Structurally, Scc1 serves as loading platform for three additional essential hook-shaped cohesin subunits called HAWKs (HEAT repeat proteins Associated With Kleisins): Scc3 interacts with kleisin constitutively, whereas Pds5 and Scc2/NIPBL compete for binding^7^. The Scc2 HAWK heterodimerizes with Scc4/MAU2 partner protein forming a so-called loader complex that promotes cohesin chromatin binding and loop extrusion^8^. In vitro, Scc2/NIPBL alone can stimulate the ATPase activity of cohesin in the presence of DNA and is sufficient for cohesin DNA loading and loop extrusion^7,9,10^. This involves clamping of DNA by Scc2 and ATP-dependent engagement of cohesin’s ATPase head domains^11,12^. Replacement of Scc2 by Pds5 on kleisin abrogates cohesin’s ATPase. Our recent results indicate that the essential role of Pds5 in budding yeast is to counteract SUMO-chain-targeted proteasomal turnover of cohesin on chromatin^13^. Of interest, this essential function of Pds5 is also bypassed by simultaneous loss of the cohesin releaser Wpl1 and the PCNA unloader Elg1^13^. In attempts to understand the mechanism underlying viability in *elg1Δ wpl1Δ pds5Δ* cells, we discovered that Scc2/NIPBL possesses a C-terminal PCNA-interacting protein (PIP) motif that recruits the cohesin loader to chromatin during replication to support de novo cohesin loading onto replicated sister DNAs and ensure SCC.

### Scc2 has a C-terminal PCNA-binding motif

We previously showed that kleisin Scc1 is targeted for proteasomal degradation by SUMO chains upon Pds5 loss^13^. Accordingly, fusing catalytically active SUMO-chain-trimming protease Ulp2 to Scc1 prevents kleisin turnover and provides viability to cells lacking Pds5^13^. To address if specifically kleisin, but not other cohesin subunits, is prone to degradation in the absence of Pds5, we asked if *GAL* promoter-mediated overexpression of *SCC1* can provide viability in *pds5Δ*. We could retrieve viable *pGAL1-SCC1 elg1Δ pds5Δ* spores upon tetrad dissection, but not *pGAL1-SCC1 pds5Δ* double mutants. Moreover, expression of *SCC1* at lower levels due to reduction of galactose concentration in the media resulted in the lethality of *pGAL1-SCC1 elg1Δ pds5Δ* cells (Extended Data Fig. 1a). Thus, the essential role of Pds5 is bypassed by loss of the PCNA unloader Elg1 when combined with either loss of the cohesin releaser Wpl1 or with overexpression of the kleisin Scc1 prone to SUMO-chain-targeted degradation in *pds5Δ* cells. Interestingly, Wpl1-mediated cohesin unloading requires Pds5 in vitro^14^ but *elg1Δ pds5Δ* cells rely on *WPL1* loss for viability^13^. This suggested that Wpl1 can unload cohesin independently of Pds5 in vivo, likely through its interaction with Scc3^15^. We therefore tested if *scc3-K404E*, defective in binding Wpl1, is able to provide viability to *elg1Δ pds5Δ* cells similar to *wpl1Δ*. This was indeed the case (Fig. 1a). Thus, increasing the chromatin-bound levels of cohesin by preventing its Wpl1-mediated unloading or by overexpressing degradation-prone kleisin Scc1 supports viability of cells lacking Pds5. Notably, these outcomes are only possible when the PCNA unloader Elg1 is additionally deleted.

**Fig. 1:**
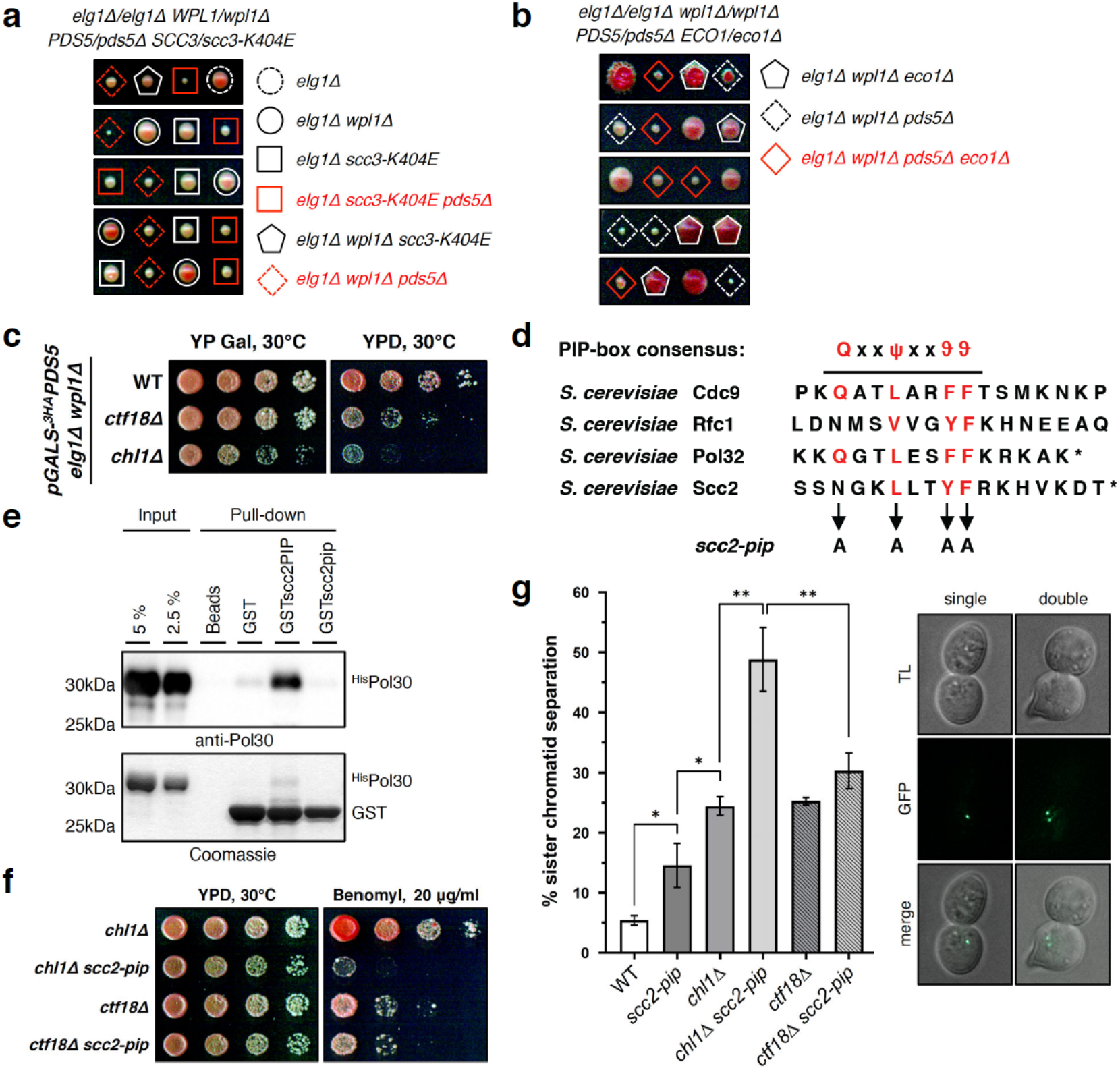
Yeast cohesin loader Scc2 harbors C-terminal PCNA-binding motif required for sister chromatid cohesion. **a**, Yeast cells lacking essential cohesin subunit Pds5 are viable in the absence of both PCNA unloader Elg1 and cohesin releaser Wpl1, or when the latter cannot bind to Scc3 cohesin subunit due to *scc3-K404E* mutation. **b**, Viability of *elg1Δ wpl1Δ pds5Δ* cells does not depend on Eco1. **c**, Growth of *elg1Δ wpl1Δ* cells upon *PDS5* shut-off relies on *CTF18* and *CHL1*. **d**, C terminus of Scc2 contains a consensus PCNA-interacting protein (PIP) motif, where ψ is any hydrophobic residue, ϑ is any aromatic residue, and x is any amino acid. Asterisk indicates the end of protein sequence. **e**, PIP of Scc2 fused to GST interacts with yeast PCNA (^His^Pol30) in vitro. **f**, *scc2-pip* mutant shows additive sensitivity to benomyl with *chl1Δ*, but is epistatic with *ctf18Δ*. **g**, *scc2-pip* mutant shows severe cohesion defects in combination with *chl1Δ*, but not with *ctf18Δ*. Data are mean values ± SD. Statistical analysis was performed using unpaired two-tailed Student’s *t*-test.

We then asked how loss of Elg1 contributes to the viability of *elg1Δ wpl1Δ pds5Δ* cells. Increased levels of DNA-loaded PCNA in the absence of Elg1 might recruit the acetyl transferase Eco1^16^ required to establish SCC by acetylating cohesin^17-21^. By dissecting diploids homozygous for *elg1Δ* and *wpl1Δ* alleles but heterozygous for *ECO1/eco1Δ* and *PDS5/pds5Δ*, we found that Eco1 was not essential for viability in *elg1Δ wpl1Δ pds5Δ* cells (Fig. 1b). Interestingly, for SCC formation during replication, the presence of the replisome-associated alternative PCNA loader Ctf18-RFC^22,23^, which participates in the de novo cohesin loading by an unknown mechanism^6^, is important^6,24^. We asked if *elg1Δ wpl1Δ pds5* cells rely on Ctf18 as well as on Chl1 helicase, the component of cohesin conversion pathway^6^, for viability. Both Ctf18 and Chl1 contributed to normal proliferation following transcriptional shut-off of *PDS5* expressed from the galactose-inducible promoter in *elg1Δ wpl1Δ* cells (Fig. 1c). Because de novo cohesin loading pathway mediated by Ctf18-RFC requires the cohesin loader Scc2^6^, we hypothesized that in *elg1Δ wpl1Δ pds5Δ* cells, the elevated chromatin-bound PCNA pool recruits Scc2 to ensure enough DNA-loaded cohesin for viability. Most PCNA-interacting proteins harbor a short sequence motif called PIP that can fit into a cavity on the surface of PCNA^25^. We in fact found a potential PIP motif at the very C terminus of Scc2 (Fig. 1d) and mutated it by replacing conserved residues with alanines to generate *scc2-pip*. In vitro GST pull-down using GST fused with the last 18 amino acids of Scc2 containing the potential PIP revealed that this peptide is indeed interacting with PCNA (yeast Pol30), whereas mutated PIP fails to do so (Fig. 1e). Thus, the cohesin loader Scc2 harbors a C-terminal PIP located within the flexible helix on the side opposite to the one with which Scc2 interacts with Smc1 and Smc3 cohesin subunits (Extended Data Fig. 1b), as judged from the recent cryo-EM structure of the budding yeast cohesin-Scc2-DNA complex^11^.

### Scc2 PIP acts in cohesin de novo loading

To study the role of Scc2 PIP in SCC, we next generated *scc2-pip* additionally carrying a C-terminal 6HA-tag. *scc2-pip* mutant variant expressed at levels similar with wild-type Scc2 (Extended Data Fig. 1c). However, when attempting to obtain *elg1Δ wpl1Δ pds5Δ SCC2-6HA* strain as control for *elg1Δ wpl1Δ pds5Δ scc2-pip-6HA*, we observed that the C-terminal tagging alone negatively affects Scc2 function, as deduced from the lethality of *elg1Δ wpl1Δ chl1Δ SCC2-6HA* (Extended Data Fig. 1d-f). Therefore, we generated *scc2-pip* mutant without epitope tagging and observed that similar to *ctf18Δ* (Fig. 1c), it causes slower proliferation and increased sensitivity to the microtubule poison benomyl in *elg1Δ wpl1Δ pds5Δ* cells (Extended Data Fig. 1g). If Scc2 is recruited via its PIP to PCNA loaded by the Ctf18-RFC in the de novo cohesin loading pathway, then one would expect *scc2-pip* mutant to have epistatic relationship with *ctf18Δ* and strong negative genetic interactions with mutants of the cohesin conversion pathway (*chl1Δ, ctf4Δ, csm3Δ, tof1Δ*). It was indeed the case when assessed by benomyl sensitivity (Fig. 1f and Extended Data Fig. h-j). Moreover, strong additive SCC defects were observed when *scc2-pip* was combined with *chl1Δ*, but not *ctf18Δ* (Fig. 1g), as measured by GFP-based cytological assay^26^. Thus, the Scc2 PIP plays a role in the de novo cohesin loading pathway of SCC.

### Scc2 PIP becomes essential without Scc4

Besides the interaction with PCNA discovered here, Scc2 is robustly localized to chromatin at centromeres via the binding of its partner Scc4 to the DDK-phosphorylated kinetochore protein Ctf19^27^. To expose the importance of Scc2 PIP for its chromatin localization we dissected *CTF18*/*ctf18Δ CTF19*/*ctf19Δ* and *CTF18*/*ctf18Δ CTF19*/*ctf19Δ scc2-pip/scc2-pip* diploid cells (Extended Data Fig. 2a,b). Similar to *ctf18Δ*, combining *scc2-pip* with *ctf19Δ* resulted in additive sensitivity to benomyl (Extended Data Fig. 2c). Furthermore, *ctf18Δ ctf19Δ scc2-pip* cells were much more slow-growing than *ctf18Δ ctf19Δ*, but viable, suggesting that Scc2 is still recruited to DNA, perhaps through Scc4-mediated binding to the chromatin remodeler RSC^28^. To completely exclude any possible Scc4-mediated chromatin localization of Scc2, we decided to use *scc2-E822K* mutant that bypasses the requirement of Scc4 for cell viability^7^, likely by providing increased binding of the mutant to DNA^11^. When we dissected *SCC2*/*scc2-E822K SCC4*/*pGALS-SCC4* diploid cells on glucose-containing plates, *scc2-E822K* suppressed the lethality of Scc4-depleted cells due to *SCC4* transcriptional shut-off (Fig. 2a). On the contrary, *scc2-E822K-17aa* mutant lacking the last 17 residues of Scc2 containing PIP (Extended Data Fig. 2d) or *scc2-E822K-pip* mutant (Fig. 2b) were no longer viable upon Scc4 loss. Thus, Scc2 PIP becomes essential in the absence of Scc4. Importantly, substituting the endogenous Scc2 PIP with the C-terminal PIP of polymerase δ nonessential subunit Pol32^29^ (Fig. 2c) provided viability to Scc4-depleted cells (Fig. 2d), but not when conserved residues of Pol32 PIP were replaced by alanines (Fig. 2e). Interestingly, when we fused Pol32 PIP downstream of mutated endogenous Scc2 PIP, it failed to support viability in *scc4* cells (Fig. 2f), suggesting that other residues of Scc2 might contribute to interaction with PCNA and that extending the C-terminus of Scc2 precludes it. Finally, instead of mutating Scc2 PIP, we combined the disassembly-prone PCNA mutant *pol30-D150E*^*30,31*^ with *pGALS-SCC4 scc2-E822K*. Upon *SCC4* shut-off, cells remained viable, but showed strong benomyl sensitivity (Extended Data Fig. 2e), further supporting the notion that the DNA-bound PCNA pool recruits Scc2 via its PIP to ensure SCC.

**Fig. 2:**
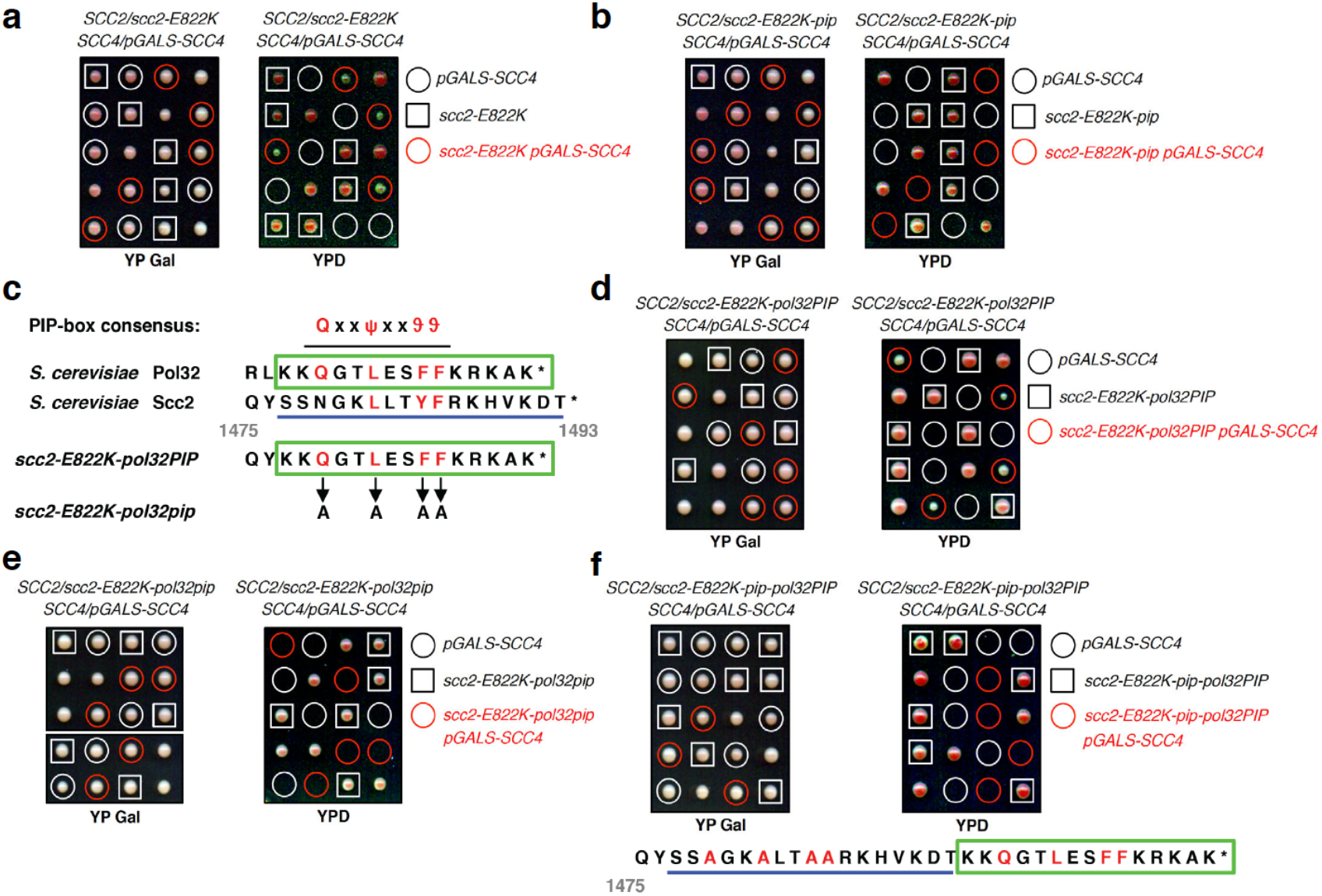
PIP motif of Scc2 becomes essential in the absence of Scc4. **a**, Tetrad dissection analysis of *SCC2*/*scc2-E822K SCC4*/*pGALS-SCC4* diploid cells. *scc2-E822K* mutant rescues the lethality of cells expressing *SCC4* from the galactose-inducible *pGALS* promoter when tetrads are dissected on glucose-containing YPD plates. **b**, *scc2-E822K-pip* mutant cannot rescue the lethality of cells upon *SCC4* shut-off. **c**, Scheme outlining the replacement of endogenous PIP of Scc2 by the C-terminal PIP of polymerase δ nonessential subunit Pol32. **d-e**, *scc2-E822K-pol32PIP* mutant (**d**), but not *scc2-E822K-pol32pip* mutant having conserved residues of Pol32 PIP replaced by alanines (**e**), is able to rescue the lethality of cells upon *SCC4* shut-off. **f**, *scc2-E822K-pip-pol32PIP* mutant carrying C-terminal fusion of Pol32 PIP downstream of mutated Scc2 PIP cannot rescue the lethality of cells upon *SCC4* shut-off.

### Scc2 PIP recruits the loader to chromatin

To evaluate the importance of Scc2 PIP for the PCNA-guided chromatin recruitment of the cohesin loader, we tagged *scc2-E822K* N-terminally with 7His8FLAG (HF)-tag and expressed it together with its various PIP mutants from either the strong constitutive promoter *pADH1* or endogenous *pSCC2* (Fig. 3a). In vitro GST pull-down using GST-PCNA precipitated ^HF^Scc2-E822K from yeast cell lysate (Extended Data Fig. 3a), whereas Ni-NTA pull-down under denaturing conditions following formaldehyde crosslinking of yeast cultures isolated cross-linked PCNA species in cells expressing ^HF^Scc2-E822K, but not ^HF^Scc2-E822K-pip (Extended Data Fig. 3b). To confirm that the isolated cross-linked species recognized by Pol30 antibody are indeed PCNA, Pol30 was C-terminally 3MYC-tagged and slower-migrating species detected by anti-MYC following crosslinking (Extended Data Fig. 3c). Interestingly, combining *POL30-3MYC* with *scc4Δ scc2-E822K* resulted in increased benomyl sensitivity (Extended Data Fig. 3d), suggesting that tagging PCNA might affect Scc2 binding. This prompted us to check mutants *pol30-6* and *pol30-79* that cause disruptions of a surface cavity on the front face of the PCNA ring^32^, which might contribute to Scc2-PCNA interaction besides Scc2 PIP. Combining *pol30-6* and *pol30-79* with *scc4Δ scc2-E822K* resulted in synthetic sickness and increased benomyl sensitivity (Extended Data Fig. 3e-g). Taken together, these results confirm physical interaction between PCNA and Scc2 mediated by the front face of the PCNA ring and Scc2 PIP.

**Fig. 3:**
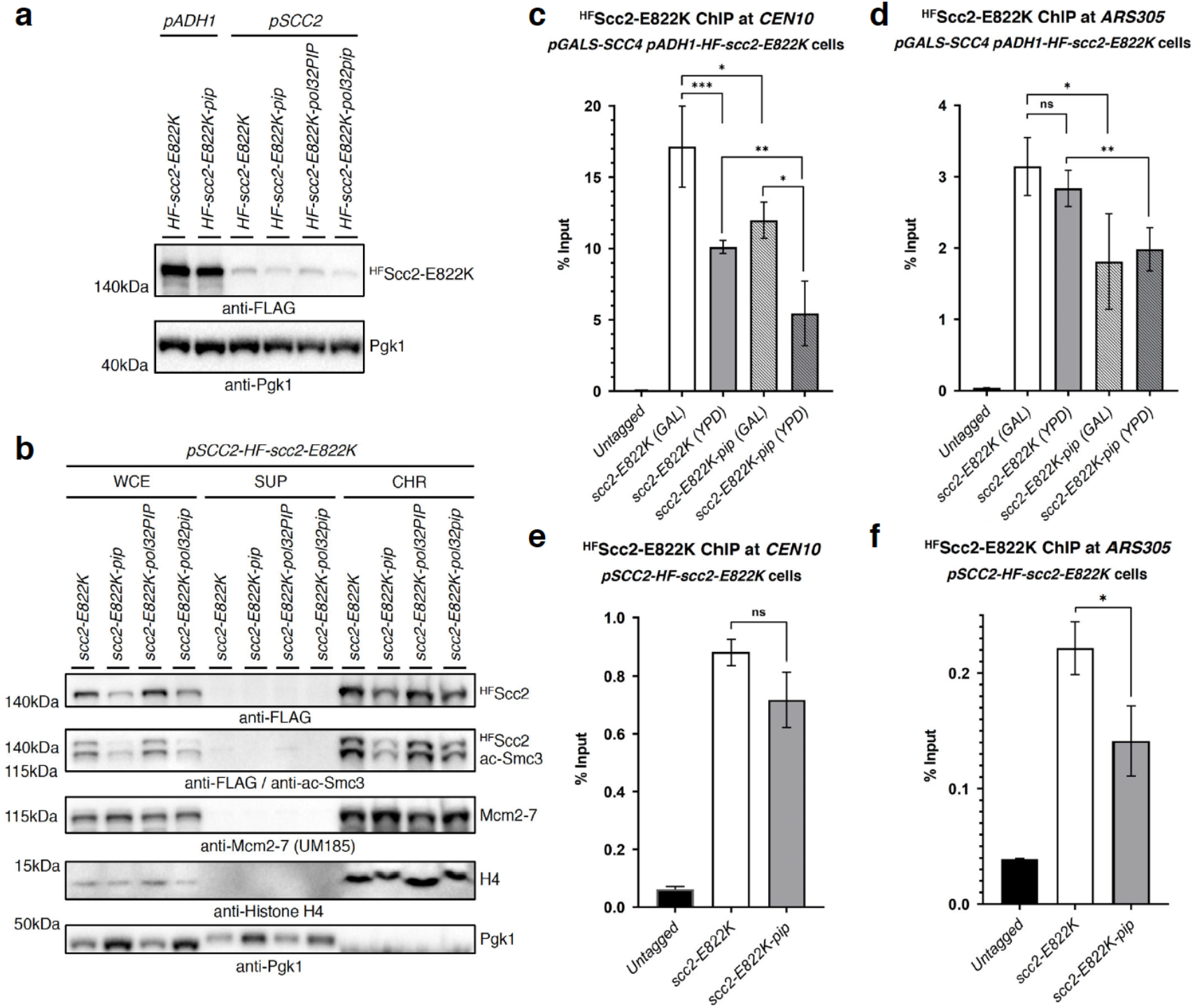
PIP motif of Scc2 ensures its proper chromatin binding. **a**, Protein levels of N-terminally 7His8FLAG (HF)-tagged Scc2-E822K and its various PIP mutants expressed from either endogenous promoter *pSCC2* or a strong constitutive promoter *pADH1*. **b**, Subcellular fractionation of cycling cells expressing HF-tagged Scc2-E822K and its various PIP mutants into soluble supernatant (SUP) and chromatin-enriched (CHR) fractions by centrifugation of the whole cell extract (WCE). Chromatin binding of ^HF^Scc2-E822K-pip is decreased compared to ^HF^Scc2-E822K and accompanied be decrease in ac-Smc3 levels. To control chromatin fractionation efficiency, the levels of histone H4, replicative helicase Mcm2-7 and the cytoplasmic/plasma membrane protein Pgk1 were detected in fractions. **c-d**, Loading of ^HF^Scc2-E822K or its PIP mutant variant expressed from strong constitutive promoter *pADH1* onto chromatin at centromere *CEN10* (**c**) and the centromere-distal early replication origin *ARS305* (**d**) was analyzed by ChIP-qPCR assay. Used cells in addition had *SCC4* expressed from galactose-inducible *pGALS* promoter allowing to study ^HF^Scc2-E822K chromatin binding upon *SCC4* shut-off after shift from galactose-containing media (GAL) to glucose (YPD). Untagged *scc2-E822K* strain was used as control. Each ChIP experiment was repeated at least three times, and each real-time PCR was performed in triplicates. Statistical analysis was performed using Student’s unpaired *t*-test. The mean values ± SD are plotted. **e-f**, ChIP-qPCR analysis as in (**c**-**d**), but ^HF^Scc2-E822K or its PIP mutant variant as well as *SCC4* are expressed from their endogenous promoters.

Next, we assayed the chromatin binding properties of ^HF^Scc2-E822K and its various PIP mutants expressed from the endogenous promoter *pSCC2* using chromatin fractionation (Fig. 3b) after having confirmed that N-terminal tagging does not change their genetic interaction with *scc4* (Extended Data Fig. 4a-d) compared to untagged *scc2* mutants (Fig. 2). We observed reduced chromatin binding of ^HF^Scc2-E822K-pip compared to ^HF^Scc2-E822K. This decrease in chromatin binding was accompanied by decreased levels of chromatin-bound acetylated Smc3, indicative of the cohesive pool of cohesin in cells. Substitution of endogenous Scc2 PIP by Pol32 PIP supported normal chromatin binding and acetylated Smc3 levels, which was not the case for mutated Pol32 PIP (Fig. 3b). Moreover, *SCC4* shut-off caused reduction in chromatin binding of ^HF^Scc2-E822K, which was additive with the decrease caused by the *scc2-pip* mutation (Extended Data Fig. 4e). Thus, both Scc4 and Scc2 PIP contribute to the chromatin binding of the cohesin loader. Finally, we quantitatively assayed ^HF^Scc2-E822K binding at the centromere *CEN10* and the early origin of replication *ARS305* by chromatin immunoprecipitation (ChIP). Upon *SCC4* shut-off (switch from GAL to YPD), the binding of ^HF^Scc2-E822K and its PIP mutant expressed from the strong *pADH1* promoter dropped specifically at centromere, but not replication origin (Fig. 3c,d). Mutation of Scc2 PIP reduced its binding at both locations. When ^HF^Scc2-E822K and its PIP mutant were expressed from the endogenous *pSCC2* promoter, their ChIP efficiency dropped tenfold at both genomic loci (Fig. 3e,f), and ^HF^Scc2-E822K-pip showed statistically significant reduction in chromatin binding at *ARS305*, but not *CEN10*. Thus, Scc2 expression levels affect the overall degree of its chromatin binding. Importantly, Scc2 PIP contributes to its recruitment at replication origins, whereas Scc4 is more important for Scc2 localization at centromeres.

### PCNA-guided Scc2 recruitment is conserved

We next asked whether the same mechanism is conserved in vertebrate cells. To this end, we performed co-immunoprecipitation experiments, revealing that NIPBL indeed interacts with PCNA in human TK6 cells as well as in chicken DT40 cells (Fig. 4a, Extended Data Fig. 5a). Based on the Scc2/NIPBL cryo-EM structures, structure predictions and motif analysis^11,12,33-35^, we found three potential PIP-like motifs at the NIPBL C-terminus resembling the Scc2 PIP (Fig. 4b,c and Extended Data Fig. 5b,c). In vitro GST pull-down assay using recombinant chicken PCNA and NIPBL C-terminal fragment (C202) confirmed the direct interaction (Fig. 4c,d). Since NIPBL PIP1 is more closely resembling the yeast Scc2 PIP among the three PIP-like motifs, we mutated PIP1 and tested the effect on its interaction with PCNA. The interaction was only moderately reduced (Fig. 4d), implying the contribution of other potential PIP-like motifs. In fact, all PIP-like motifs were able to interact with PCNA in vitro, and the interaction was weakened when they were mutated (Extended Data Fig. 5d).

**Fig. 4:**
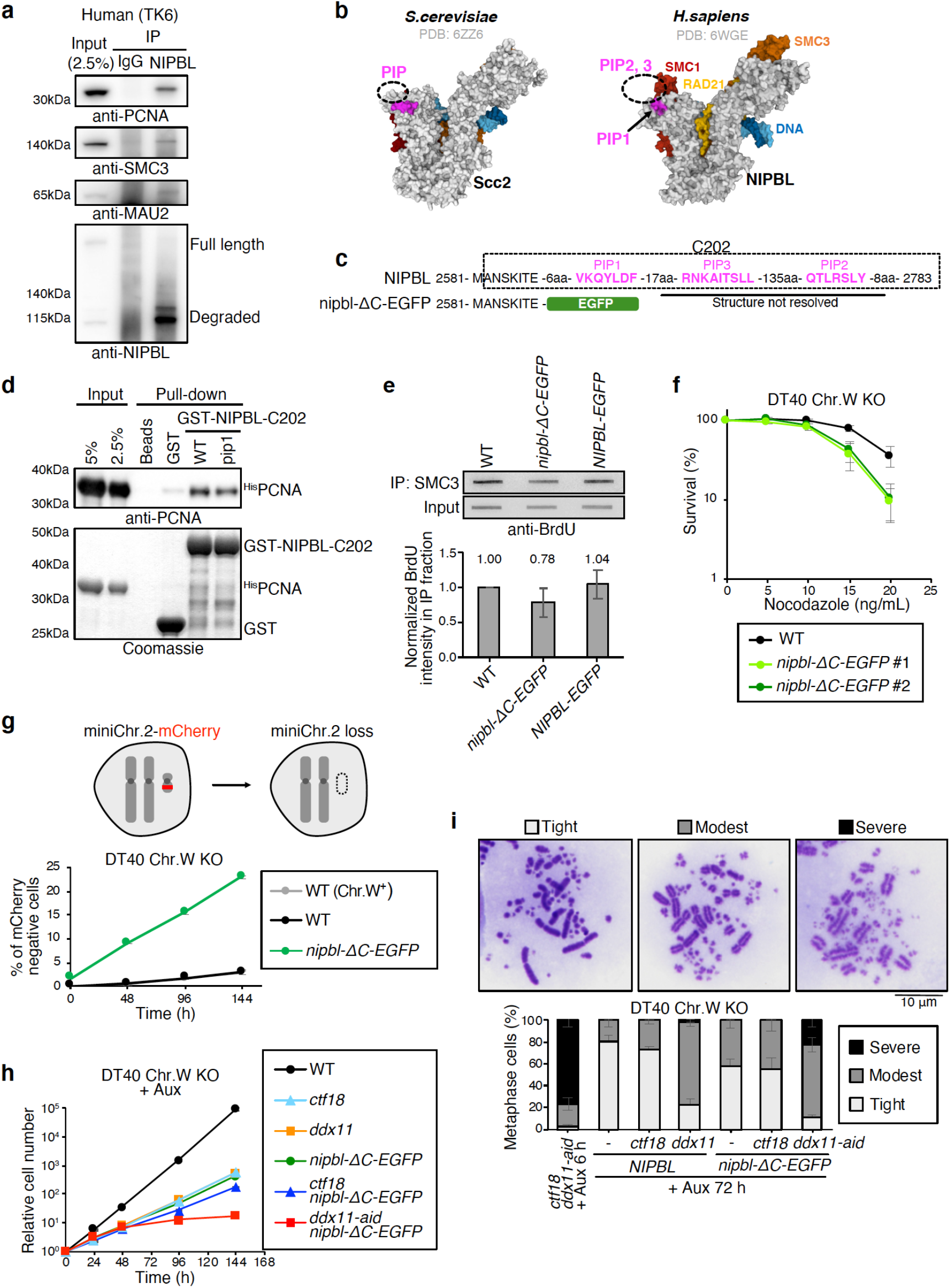
PCNA-guided recruitment of chicken cohesin loader NIPBL to chromatin is required for sister chromatid cohesion. **a**, Immunoprecipitation of human NIPBL in TK6 cells. PCNA was co-immunoprecipitated with NIPBL. SMC3 and MAU2 are shown as positive control. **b**, Cryo-EM structures of budding yeast Scc2-cohesin (PDB: 6ZZ6) and human NIPBL-cohesin (PDB: 6WGE). The DNA is colored blue, SMC1 – red, SMC3 – orange, RAD21 – yellow. 10 amino acids (FSAQLENIEQ) upstream of budding yeast Scc2 PIP, and PIP1 of human NIPBL are colored pink. Dashed circles indicate where unmapped PIPs should be positioned. **c**, C-terminus of chicken NIPBL containing three PIP-like motifs. *nipbl-ΔC-EGFP* cells lack last 195 amino acids, and EGFP-tag is fused instead. **d**, C-terminal fragment of NIPBL (C202) fused to GST interacts with chicken PCNA in vitro. Mutating PIP1 weakened the interaction. **e**, The amount of SMC3 on newly replicated chromatin was determined by BrdU-ChIP-Slot-Western technique. Truncation of the last 195 aa of NIPBL results in the reduction of SMC3 levels bound to nascent DNA. The mean values of 5 independent experiments ± SE are plotted. **f**, Sensitivity assay using CellTiter-Glo. *nipbl-ΔC-EGFP* mutant exhibits hypersensitivity to nocodazole treatment. The mean values of three independent experiments ± SE are plotted. **g**, Frequency of chromosome loss was measured by mini-chromosome loss assay. Mini-chromosome 2 carries mCherry expression unit, and the percentage of mCherry negative cells was determined by flow cytometry. The mean values of three independent experiments ± SD are plotted. **h**, Growth curves showing synthetic lethality of *nipbl-ΔC-EGFP* with *ddx11*, but not with *ctf18*. **i**, Cohesion analysis of metaphase spreads. Depletion of DDX11 in *nipbl-ΔC-EGFP* background results in severe cohesion defects.

Next, to assess the consequence of the defect in PCNA-NIPBL interaction, we established *nipbl-ΔC-EGFP* cells, in which the last 195 residues of NIPBL were replaced with EGFP (Fig. 4c). Because chicken *NIPBL* genes are located on chromosome Z and W, we utilized DT40 cells lacking chromosome W, which carry a single copy of *NIPBL* (Extended Data Fig. 5e). Subsequently, the EGFP-tag was introduced to the C-terminus of NIPBL using the FLP-In system, with or without truncating the last 195 residues where the three PIP-like motifs are positioned (Extended Data Fig. 5f-h). We then employed BrdU-ChIP-Slot-Western technique^36^ to monitor the cohesin recruitment to newly replicated chromatin. Notably, the amount of BrdU co-immunoprecipitated with SMC3 was reduced in *nipbl-ΔC-EGFP* cells, suggesting that the PCNA-NIPBL interaction facilitates cohesin loading on nascent chromatin (Fig. 4e). Moreover, *nipbl-ΔC-EGFP* cells exhibited hyper-sensitivity to nocodazole (Fig. 4f, Extended Data Fig. 5i) and increased chromosome loss rate (Fig. 4g), measured by the mini-chromosome loss assay^37^. Thus, these results confirm the importance of PCNA-guided NIPBL recruitment for proper SCC.

*scc2-pip* and mutants of the de novo cohesin loading pathway factors showed epistatic genetic interaction in budding yeast (Fig. 1f-g). To test if the same holds in vertebrates, we aimed to knockout *CTF18* and *DDX11* in *nipbl-ΔC-EGFP*. Notably, we could disrupt *CTF18*, but not *DDX11* (Extended Data Fig. 5j). Therefore, we employed the auxin inducible degron (AID) system to obtain *ddx11-aid* conditional mutant^38,39^ (Extended Data Fig. 5k). Remarkably, *ddx11-aid nipbl-ΔC-EGFP* cells stopped proliferating 4 days after auxin addition (Fig. 4h), indicative of synthetic lethality. Moreover, strong additive SCC defects were observed when *nipbl-ΔC-EGFP* was combined with *ddx11*, but not with *ctf18* (Fig. 4i), consistent with the findings in yeast (Fig. 1g). Taken together, we conclude that PCNA-guided Scc2/NIPBL recruitment onto replicated DNA is a fundamental mechanism to ensure SCC, conserved from yeast to vertebrates.

In summary, our study uncovers a conserved mechanism of de novo cohesin loading during DNA replication necessary for SCC. This mechanism relies on the recruitment of the cohesin loader Scc2/NIPBL to the PCNA sliding clamp, deposited onto newly-replicated DNA by the alternative PCNA loader CTF18-RFC associated with the replisome. Previously, cohesin and NIPBL loaded at replication origins were proposed to remain associated with the replicative helicase MCM and then transferred behind the replication fork to establish SCC^40^. Our work suggests that PCNA, the maestro of various replication-linked functions, serves as a recruiting platform for incoming Scc2/NIPBL. Thus, PCNA coordinates Scc2/NIPBL-mediated cohesin de novo loading unto replicated DNA with cohesin acetylation required for SCC establishment and mediated by Eco1/ESCO2 acetyltransferase^17-21^, likewise recruited by the homotrimeric PCNA ring^41,42^.

## Methods

### Yeast strains, techniques and growing conditions

Chromosomally tagged *Saccharomyces cerevisiae* strains and mutants were constructed by a PCR-based strategy, by genetic crosses and standard techniques^43^. Standard cloning and site-directed mutagenesis techniques were used. Strains and all genetic manipulations were verified by polymerase chain reaction (PCR), sequencing and phenotype. Maps and primer DNA sequences are available upon request. All yeast strains used in this work are isogenic to W303 background and are listed in the Supplementary Table 1. Yeast cultures were inoculated from overnight cultures, grown using standard growth conditions and media^44^. All cultures were grown in YPD media containing glucose (2%) as carbon source at 30°C unless otherwise indicated. For the transcriptional shut-off of genes expressed under the control of *GAL* promoter, cells were grown in YP Gal media containing galactose (2%), washed once with 1X PBS and shifted to YPD media or plated on YPD plates. For cell cycle synchronization, logarithmic cells grown at 30°C were arrested in G1 using 3-5 μg/ml of alpha-factor for 2-3 hours. G2/M arrest was performed with 20 μg/ml of nocodazole for 2-3 hours. G1/G2-arrest was verified microscopically and by FACS analysis. For drug sensitivity assays, cells from overnight cultures were counted and diluted before being spotted on YPD plates containing the indicated concentrations of benomyl and incubated at 30°C for 2-3 days. The tetrad dissection analysis was performed using the Singer Instruments MSM 400 system on YPD or YP Gal plates.

### Premature sister chromatid separation assay in budding yeast

Sister chromatid cohesion was measured as described previously^26^. Logarithmically growing cells were treated with 3 μg/ml alpha-factor to induce G1 arrest. Cells were then washed using YP and released in YPD containing 20 μg/ml of nocodazole in order to allow one round of replication. After three hours of nocodazole treatment, G2/M arrest was checked by cell morphology and cells were collected, washed once with 1X PBS and fixed in 70% ethanol overnight at - 20°C. Cells were then resuspended in 50 mM Tris HCl, pH 6.8 and sonicated for 5 seconds prior to microscopic analysis. Cells were imaged on a DeltaVision microscope (Applied Precision) using 100X oil immersion lens. Images were analyzed using ImageJ software. Statistical analysis was performed on results obtained in at least three independent experiments using Student’s unpaired *t*-test, at least 240 cells were analyzed per each strain. The error bars represent standard deviation (SD).

### GST in vitro pull-down assays

The C-terminus of yeast Scc2 (aa 1476-1493) either wild-type (termed GSTscc2PIP) or with PIP mutated (Scc2-N1479A,L1482A,Y1485A,F1486A, termed GSTscc2pip) was introduced into pGEX-6P-1 (GE Healthcare) vector replacing the last 18 residues of GST. The C-terminus of chicken NIPBL (aa 2582-2783) either wild-type (termed GST-NIPBL WT) or with PIP1 mutated (NIPBL-Q2597A,L2599A,F2601A, termed GST-NIPBL pip1) was introduced into pGEX-6P-1 vector replacing the last 13 residues of GST. In addition, sequences containing individual potential PIP motifs of the chicken NIPBL C-terminus (aa 2586-2608, termed GST-NIPBL-PIP1; aa 2764-2783, termed GST-NIPBL-PIP2; aa 2616-2635, termed GST-NIPBL-PIP3) or having their conserved residues mutated (NIPBL-Q2597A,L2599A,F2601A, termed GST-NIPBL-pip1; NIPBL-Q2769A,L2771A,Y2775A, termed GST-NIPBL-pip2; NIPBL-R2626A,N2627A,I2630A,L2633A,L2634A, termed GST-NIPBL-pip3) were introduced into pGEX-6P-1 vector replacing the last 13 residues of GST. Rosetta (DE3) pLysS competent *E. coli* cells (Novagen) were used for protein expression. Following overnight protein induction with 0.25 mM IPTG at 16ºC in 100 ml cell cultures, cells were pelleted, resuspended in 6 ml lysis buffer (1X PBS, 500 mM NaCl, 1% Triton X-100, lysozyme, Calbiochem EDTA-free protease inhibitor cocktail set III) and sonicated on ice. The crude lysate was clarified by centrifugation at 15000 rpm for 15 min at 4°C, and the supernatant was mixed with 0.2 ml of glutathione sepharose 4B beads (GE Healthcare) pre-equilibrated with lysis buffer. Following overnight incubation at 4°C, five washes with lysis buffer were performed and the beads with bound GST-fusion proteins were used for subsequent in vitro pull-down assays with recombinant PCNA (either yeast N-terminally His-tagged Pol30 with yeast Scc2 GST fusions or chicken N-terminally His-tagged PCNA with chicken NIPBL GST fusions). The amounts of GST-fusion proteins bound to the beads were estimated by comparison to the BSA samples of known concentrations resolved by SDS-PAGE and Coomassie Blue staining. To study the interaction between GSTscc2PIP fusions and PCNA, purified recombinant His-tagged yeast Pol30 (2.5 μg) was incubated either with GSTscc2PIP (2.5 μg) and its PIP mutant variant GSTscc2pip, or GST (2.5 μg) alone bound to glutathione sepharose 4B beads in 0.7 ml of binding buffer (1X PBS, 150 mM NaCl, 1% Triton X-100, Calbiochem EDTA-free protease inhibitor cocktail set III) overnight at 4°C, with gentle mixing. After the incubation, the beads were washed 5 times with 1 ml of the binding buffer and bound proteins were eluted with 50 μl of HU sample buffer. Samples were then analyzed by SDS-PAGE followed by Western blotting and probing with anti-Pol30 (GTX64144, Gene Tex), and in parallel by staining of the protein gel with Coomassie Blue. To study the interaction between full-length Scc2 and PCNA, recombinant GST-Pol30 fusion was purified and used for GST pull-down from whole cell extracts obtained by grinding in liquid nitrogen of yeast cells expressing N-terminally 7His8FLAG-tagged Scc2-E822K (^HF^Scc2-E822K) from a strong constitutive *pADH1* promoter. Anti-PCNA (sc-25280, Santa Cruz Biotechnology) and anti-GST antibodies were used to study the interaction between chicken PCNA and GST-NIPBL fusions.

### TCA protein precipitation

For preparation of denatured protein extracts, yeast cultures grown to an OD_600_=0.7-1 were pelleted by centrifugation (4000 rpm, 4 min, 4°C) and immediately frozen in liquid nitrogen. After thawing on ice, the pellets were lysed by addition of denaturing lysis buffer (1.85 M NaOH, 7.5% β-mercaptoethanol) for 15min on ice. For the cell pellet of an OD_600_=1 typically 150 μl of lysis buffer was used. To precipitate the proteins, the lysate was subsequently mixed with an equal volume (150 μl in case of OD_600_=1) of 55% (w/v) trichloroacetic acid (TCA) and further incubated on ice for 15 min. The precipitated material was recovered by two sequential centrifugation steps (13000 rpm, 4°C, 15 min). Pelleted denatured proteins were then directly resuspended in HU sample buffer (8 M urea, 5% SDS, 1 mM EDTA, 1,5% DTT, 1% bromophenol blue; 50 μl per OD_600_=1), boiled for 10 min and stored at −20°C. Proteins were resolved on precast Bolt 4%–12% Bis-Tris Plus gradient gels, and analyzed by standard Western blotting techniques. Mouse monoclonal anti-FLAG antibody (1:2000, clone M2) was purchased from Sigma-Aldrich. Mouse monoclonal anti-Pgk1 antibody (1:2000; clone 22C5D8) was obtained from Thermo Fisher Scientific. Mouse monoclonal anti-HA (1:2000; clone F-7) and anti-PCNA (1:2000; clone F-2) antibodies were from Santa Cruz Biotechnology, as well as normal mouse IgG. Rabbit polyclonal anti-Pol30 antibody (1:2000; GTX64144) was purchased from Gene Tex. Mouse monoclonal anti-c-MYC antibody (1:2000; clone 9E10) and rabbit polyclonal anti-GST antibody (1:2000) were produced in house. Rabbit polyclonal anti-Histone H4 antibody (1:2000; ab7311) was obtained from Abcam. Mouse monoclonal anti-acetyl-Smc3 antibody^45^ (1:2000) was a gift from Katsuhiko Shirahige. Rabbit polyclonal anti-Mcm2-7 antibody^46^ (1:5000; UM185) was a gift from Stephen P. Bell. Anti-rabbit IgG and anti-mouse IgG, HRP-linked antibodies (1:5000) were purchased from Cell Signaling Technology.

### Ni-NTA pull-down of 7His8FLAG-tagged Scc2-E822K after formaldehyde crosslinking of yeast cells

For isolation of Scc2 protein interactors from yeast cells expressing N-terminally 7His8FLAG-tagged Scc2-E822K (^HF^Scc2-E822K) or its PIP mutant variant, denatured protein extracts were prepared following formaldehyde crosslinking and Ni-NTA chromatography was carried out as described previously^47,48^. Briefly, 200 OD_600_=1 of logarithmically growing cells were harvested by centrifugation (4000 rpm, 4 min, 4°C) after 30 min of crosslinking with 1% formaldehyde, washed with pre-chilled water, transferred to 50 ml falcon tube and lysed with 6 ml of 1.85 M NaOH / 7.5% β-mercaptoethanol for 15 min on ice. The proteins were precipitated by adding 6 ml of 55% TCA and another 15 min incubation on ice (TCA-precipitation, described above). Next, the precipitate was pelleted by centrifugation (3500 rpm, 15 min, 4°C), washed twice with water and finally resuspended in buffer A (6 M guanidine hydrochloride, 100 mM NaH_2_PO_4_, 10 mM Tris-HCl, pH 8.0, 20 mM imidazole) containing 0.05% Tween-20. After incubation for 1 hour on a roller at room temperature with subsequent removal of insoluble aggregates by centrifugation (23000 g, 20 min, 4°C), the protein solution was incubated overnight at 4°C with 50 μl of Ni-NTA agarose beads in the presence of 20 mM imidazole. After incubation, the beads were washed three times with buffer A containing 0.05% Tween-20 and five times with buffer B (8 M urea, 100 mM NaH_2_PO_4_, 10 mM Tris-HCl, pH 6.3) with 0.05% Tween-20. ^HF^Scc2-E822K and its cross-linked species bound to the beads were finally eluted by incubation with 50 μl of HU sample buffer for 10 min at 65°C. Proteins were resolved on precast Bolt 4%–12% Bis-Tris Plus gradient gels, and analyzed by standard Western blotting techniques.

### ChIP-qPCR

Chromatin immunoprecipitation (ChIP) was carried out as previously described^49^. Briefly, cells were collected at the indicated experimental conditions and crosslinked with 1% formaldehyde for 30 min. Cells were washed twice with ice-cold 1X TBS, suspended in lysis buffer supplemented with 1 mM PMSF, 20 mM NEM, and 1X EDTA-free complete cocktail, and lysed using FastPrep-24 (MP Biomedicals). Chromatin was sheared to a size of 300-500 bp by sonication. IP reactions with anti-FLAG antibodies and Dynabeads protein G were allowed to proceed overnight at 4°C. After washing and eluting the ChIP fractions from beads, crosslinks were reversed at 65°C overnight for both Input and IP. After proteinase K treatment, DNA was extracted twice by phenol/chlorophorm/isoamyl alcohol (25:24:1, v/v). Following precipitation with ethanol and Ribonuclease A (RNase A) treatment, DNA was purified using QIAquick PCR purification kit. Real-time PCR was performed using QuantiFast SYBR Green PCR kit according to the manufacturer’s instructions and each reaction was performed in triplicates using a Roche LightCycler 96 system. The results were analyzed with absolute quantification/2^nd^ derivative maximum and the 2(-ΔC(t)) method. Each ChIP experiment was repeated at least three times. Statistical analysis was performed using Student’s unpaired *t*-test. The error bars represent standard deviation (SD).

### Chromatin fractionation

The chromatin binding assay was performed as described previously^13,47^. Briefly, native yeast protein extract was prepared from 50 OD_600_=1 of logarithmically growing culture by treating harvested cells with zymolyase to produce spheroplasts and disrupting them with 1% Triton X-100. The resulting whole cell extract (WCE) was carefully applied on top of the 30% sucrose cushion of equal volume and centrifuged for 30 min at 20000 g at 4°C. The supernatant containing soluble protein fraction (SUP) was carefully collected from the top of the cushion, sucrose aspirated and the pellet containing the chromatin fraction (CHR) was resuspended in HU sample buffer for subsequent SDS-PAGE and Western blot analysis.

### Cell lines and general techniques

Cell lines used in this study are listed in Supplementary Table 2. mRNA isolation, reverse transcription PCR, western blotting, cohesion analysis were performed as previously described^50^.

### Cell culturing

TK6 cells were cultured at 37ºC in RPMI-1640 medium supplemented with 10% horse serum, 2 mM L-glutamine, Penicillin/Streptomycin mix, 1.8 mM sodium pyruvate. DT40 cells were cultured at 39.5ºC in D-MEM/F-12 GlutaMAX supplement medium supplemented with 10% fetal bovine serum, 2% chicken serum, Penicillin/Streptomycin mix, and 10 μM 2-mercaptoethanol in the presence or absence of 500 μM Auxin. To plot growth curves, each cell line was cultured in three different wells of 24-well plates and passaged every 24 h. Cell number was determined at each time point by flow cytometry.

### Plasmid construction and transfection

To generate the chicken *NIPBL* expression construct, 3xHA-tag was inserted into EcoRV locus of pBACT-Puro vector^51^. Subsequently, full-length *NIPBL* cDNA was amplified using primers 5’-GCATgcggccgcTAATGGGGATATGCCTCATGTTCC-3’ (NotI) and 5’-GCATggtaccTTAGCTCGAAGTTCCATCCTTGG-3’ (KpnI), and cloned into the pBACT-3xHA-Puro vector. The construct was linearized by FspI before transfection. For engineering mini-chromosome 2, the *GFP* cassette of telomere seeding vector^37^ was replaced with *mCherry* cassette. Otherwise, the plasmids used for mini-chromosome loss assay and the transfection method are previously described^37^. To generate the *NIPBLw-EGFP* knock-in construct, the homology arm was amplified using primers 5’-AAAgtcgacTGCTGGATAGCGAAGATGGAGAAG-3’ (SalI) and 5’-AAAgcggccgcTTCTGCTGGTGCAGATTTCTGTG-3’ (NotI), and ligated into pLoxP vector^52^. Subsequently, *Puro-GFP* cassette was cloned into endogenous BamHI site in the middle of homology arm. The construct was linearized by NotI prior to transfection. *SMAD7-Ecogpt* knock-in construct was made by cloning the homology arm amplified by primers 5’-AAAgtcgacCTTAGGGATGGAGTGGGGCATCCAG-3’ (SalI) and 5’-AAAgcggccgcCCATCATGTCATTGGGTGCTTAGG-3’ (NotI) into pLoxP vector. *Ecogpt* cassette was then cloned into endogenous BamHI site. The construct was linearized by NotI before transfection. To truncate last 195 amino acids of NIPBL on chromosome Z and fuse EGFP, 2.2 kb of homology arm downstream of E2588 was amplified by using primers 5’-GTACgtcgacCAAACGCAGGAAGAGCCACTG-3’ (SalI) and 5’-GTACgctagcTTCAGTGATCTTGGAATTAGCCATATCAC-3’ (NheI), and cloned into pEGFP-cFLP-Eco vector^51^. The plasmid was linearized by AflII prior to transfection. For adding EGFP at the C-terminus of NIPBL, 2 kb of homology arm upstream of stop codon was amplified by primers 5’-GCATgtcgacCAGGAAGACAGGAGTGCATTTCCATC-3’ (SalI) and 5’-GCATactagtGCTCGAAGTTCCATCCTTGGC-3’ (SpeI), and cloned into pEGFP-cFLP-Eco vector. The plasmid was linearized by NheI prior to transfection.

### Drug sensitivity assay

To assess the drug sensitivity, 1×10^4^ cells were cultured in 24-well plates containing various concentrations of DNA damaging agents in 1 ml of medium in duplicates. Cell viability was assessed after 48 h by CellTiter-Glo assay following manufacturer’s protocol. Percentage of survival was determined by considering the luminescent intensity of untreated cells as 100%.

### Co-immunoprecipitation in TK6 and DT40 cells

For immunoprecipitating NIPBL in TK6 cells, 1×10^7^ cells were lysed in 0.5 mL of lysis buffer (20 mM Tris-HCl pH 7.4, 150 mM NaCl, 5 mM MgCl_2_, 0.5% NP-40, 10% Glycerol, 20 mM N-ethylmaleimide, 1 mM PMSF, 1x cOmplete, 50 U/mL benzonase). Lysates were then rotated at 4°C for 1 h, and 37°C for 10 min. After centrifugation (14000 rpm, 4°C, 20 min), supernatants were incubated with 1 μg of anti-NIPBL antibody (A301-779A, Bethyl) and 3 mg of Dynabeads Protein A for 2 h. Beads were then washed with 1 mL of wash buffer (20 mM Tris-HCl pH 7.4, 200 mM NaCl, 5 mM MgCl_2_, 0.5% NP-40, 10% Glycerol, 20 mM N-ethylmaleimide, 1 mM PMSF, 1x cOmplete) 4 times and incubated with 30 μL of 1x Laemmli buffer at 95°C for 15 min for elution. To immunoprecipitate (IP) 3HA-NIPBL in DT40 cells, 1×10^7^ cells were lysed in 1 mL of lysis buffer (20 mM Tris-HCl pH 7.4, 150 mM NaCl, 5 mM MgCl_2_, 0.5% NP-40, 10% Glycerol, 20 mM N-ethylmaleimide, 1 mM PMSF, 1x cOmplete). Following 10 min of incubation on ice, lysates were sonicated (10%, 12 sec, 3 cycles) to solubilize the chromatin. After centrifugation (14000 rpm, 4°C, 5 min) supernatants were incubated with 50 μL of Pierce Anti-HA Magnetic Beads overnight. Beads were then washed with 1 mL of wash buffer (20 mM Tris-HCl pH 7.4, 200 mM NaCl, 5 mM MgCl_2_, 0.5% NP-40, 10% Glycerol, 20 mM N-ethylmaleimide, 1 mM PMSF, 1x cOmplete) 4 times and incubated with 30 μL of 1x Laemmli buffer at 95°C for 15 min for elution. IP samples were analyzed by standard Western blotting techniques. Antibodies used were the following: anti-HA (1:1000, 12158167001, Roche), anti-GAPDH (1:1000, sc-47724, Santa Cruz Biotechnology), anti-GFP (1:500, sc-9996, Santa Cruz Biotechnology), anti-MAU2 (1:1000, ab183033, Abcam), anti-miniAID (1:1000, M214-3, MBL), anti-NIPBL (1:1000, A301-779A, Bethyl), anti-PCNA (1:2000, sc-25280, Santa Cruz Biotechnology), anti-SMC3 (1:1000, kind gift from Ana Losada).

### Mini-chromosome loss assay

Mini-chromosome 2 was engineered as previously described^37^ with small modifications. The *GFP* expression unit of telomere-seeding vector targeting *TPK1* was replaced with *mCherry* expression unit. Cells were cultured in the medium containing L-histidinol (1 mg/mL) and puromycin (0.5 μg/mL) for 48 h prior to starting the experiment in order to exclude cells that already lost mini-chromosome 2. Percentage of mCherry negative cells was determined by flow cytometry every 48 h. 10000 cells were analyzed for each condition.

### BrdU-ChIP-Slot-Western technique

BrdU-ChIP-Slot-Western technique was performed as previously described^36^ with minor modifications. 1×10^7^ cells were pulsed-labelled with 20 μM BrdU for 20 min, and cross-linked by 1% paraformaldehyde for 10 min at room temperature. Cross-link was then quenched by 125 mM glycine for 5 min at room temperature. Following centrifugation (900 g, 4°C, 10 min), pellets were washed with 10 mL of cold PBS twice. Subsequently, pellets were resuspended with 0.5 mL of cold FA140 buffer (50 mM HEPES-KOH pH 7.5, 140 mM NaCl, 1 mM EDTA pH 8.0, 1% Triton-X100, 0.1% sodium deoxycholate, 1 mM PMSF, 1x cOmplete) and sonicated to shear the chromatin (Level 6, 10 sec, 6 cycles). Following sonication, the lysates were centrifugated (14000 rpm, 4°C, 10 min). Supernatants were then incubated with 1 μg of anti-SMC3 antibody (a gift from Ana Losada) and 1 mg of Dynabeads Protein A at 4°C overnight. 0.1 mL of supernatant was taken as “Input” and kept on ice. Beads were washed by 0.5 mL of cold FA140 buffer for 5 min, and by 0.5 mL of cold FA500 buffer (50 mM HEPES-KOH pH 7.5, 500 mM NaCl, 1 mM EDTA pH 8.0, 1% Triton-X100, 0.1% sodium deoxycholate, 1 mM PMSF, 1x cOmplete) for 5 min. Subsequently, beads were washed with 0.5 mL of cold LiCl buffer (10 mM Tris-HCl pH 8.0, 250 mM LiCl, 0.5% NP-40, 0.5% sodium deoxycholate, 1 mM PMSF, 1x cOmplete). Following the wash with LiCl buffer, beads were incubated with 0.2 mL of elution buffer (1% SDS, 100 mM NaHCO_3_) at 25°C for 15 min twice for elution and collected together. 0.3 mL of elution buffer was added to “Input” samples. 13.2 μL of 5M NaCl was added to the eluted “IP” samples and “Input”, and incubated at 65°C for 5 h to reverse the crosslinks. 1 mL of ethanol was added, kept at −80°C overnight and finally centrifugated (14000 rpm, 4°C, 15 min). Pellets were washed with 0.8 mL of 70% ethanol once, and air-dried at 37°C for 10 min. Pellets were then resuspended with 90 μL of milli-Q water, and 2 μL of 10 mg/mL RNaseA was added and incubated at 37°C for 30 min. After RNaseA treatment, 10 μL of 10x Proteinase K buffer (100 mM Tris-HCl pH 8.0, 50 mM EDTA pH 8.0, 5% SDS) and 1 μL of 20 mg/mL Proteinase K were added, then incubated at 42°C for 1 h. Subsequently, DNA was purified using PCR purification kit (Qiagen) following manufacturer’s protocol, eluted by 50 μL of milli-Q water. DNA concentration was measured using Qubit dsDNA HS assay kit (Invitrogen), and the concentration was adjusted to 30 ng/μL for “Input”, and 0.25 ng/μL for IP fractions at the volume of 50 μL. Samples were then denatured by adding 125 μL of 0.4N NaOH and incubated for 30 min at room temperature. Following denaturation, 175 μL of 1 M Tris-HCl pH 6.8 was added to neutralize and samples were kept on ice. Denatured DNAs were then blotted onto nitro-cellulose membrane pre-wet by 20x SSC, and crosslinked using UV chamber (125 mJ, BioRad). Membrane was then subjected for antibody reaction following the same procedure of Western blotting. Mouse monoclonal anti-BrdU antibody (1:500, 347580, BD Biosciences) was used for BrdU detection.

## Reporting summary

Further information on research design is available in the Nature Research Reporting Summary linked to this paper.

## Data availability

The authors declare that the data supporting the findings of this study are available within the Article. Any other data related to this study are available from the corresponding authors D.B. and I.P. upon request. All data are archived at the IFOM, the Foundation Institute of Molecular Oncology or the Department of Chemistry at Tokyo Metropolitan University.

## Acknowledgments

We thank S.P. Bell, S. Jentsch, A. Losada, K. A. Nasmyth, and K. Shirahige for sharing reagents, S. Barozzi, F. Casagrande, and M. Garre for help with microscopy. This work was supported by the Italian Association for Cancer Research (IG 18976, IG 23710), and European Research Council (Consolidator Grant 682190) grants to D.B., EMBO long-term fellowship (ALTF 561-2014) and an AIRC/Marie Curie Actions - COFUND iCARE fellowship to I.P. R.K was partly supported by an AIRC 3-yr fellowship Mari e Valeria Rindi (Rif.22403). T.A. received support from Japanese Society for the Promotion of Science KAKENHI (17K17986).

## Author contributions

D.B., R.K., and I.P. conceived the study. R.K., I.P., and T.A. performed experiments. R.K., I.P., T.A., K.H., and D.B. analyzed the data. R.K. and I.P. constructed the figures. D.B., R.K., and I.P. wrote the manuscript.

## Supplementary information

Supplementary Information is available for this paper.

**Supplementary Table 1.** *Saccharomyces cerevisiae* strains used in this study.

| Strain | Relevant Genotype | Reference |
| --- | --- | --- |
| FY1363<br>(W303<br>WT) | *MATa ade2-1 can1-100 his3-11,15 leu2-3,112 trp1-1 ura3-1 RAD5+* | Lab<br>collection |
| FY1364<br>(W303<br>WT) | *MATalpha ade2-1 can1-100 his3-11,15 leu2-3,112 trp1-1 ura3-1*<br>*RAD5+* | Lab<br>collection |
| HY10598 | *MATa/MATalpha elg1::HIS3MX6/elg1::HIS3MX6*<br>*wpl1::TRP1/wpl1::TRP1 PDS5/pds5::NatMX4* | This study |
| HY10876 | *MATa/MATalpha elg1::HIS3MX6/elg1::HIS3MX6*<br>*wpl1::TRP1/wpl1::TRP1 PDS5/pds5::NatMX4*<br>*ECO1/eco1::KANMX4* | This study |
| HY10890 | *MATa/MATalpha elg1::HIS3MX6/elg1::HIS3MX6*<br>*wpl1::TRP1/wpl1::TRP1 PDS5/pds5::NatMX4 RAD51/rad51::LEU2*<br>*SGS1/sgs1::KANMX4* | This study |
| HY11326 | *MATa/MATalpha ELG1/elg1::HIS3MX6 WPL1/wpl1::TRP1*<br>*ECO1/eco1::KANMX4 SCC3/scc3-K404E-6HA::HPHNT1* | This study |
| HY11421 | *MATa/MATalpha elg1::HIS3MX6/elg1::HIS3MX6*<br>*WPL1/wpl1::TRP1 PDS5/pds5::NatMX4 SCC3/scc3-K404E-*<br>*6HA::HPHNT1* | This study |
| HY10996 | *MATa/MATalpha ELG1/elg1::HPHMX6 CHL1/chl1::KANMX4* | This study |
| HY10997 | *MATa/MATalpha ELG1/elg1::HPHMX6 CTF18/ctf18::TRP1* | This study |
| HY11097 | *MATa/MATalpha wpl1::HIS3MX6/wpl1::HIS3MX6*<br>*elg1::HPHMX6/elg1::HPHMX6 CTF18/ctf18::TRP1*<br>*CHL1/chl1::KANMX4* | This study |
| HY11053 | *MATa elg1::HIS3MX6 wpl1::TRP1 (natNT2)pGALS-3HA-PDS5* | This study |
| HY11081 | *MATa elg1::HIS3MX6 wpl1::TRP1 (natNT2)pGALS-3HA-PDS5*<br>*ctf18::KANMX4* | This study |
| HY11083 | *MATa elg1::HIS3MX6 wpl1::TRP1 (natNT2)pGALS-3HA-PDS5*<br>*chl1::KANMX4* | This study |
| HY11104 | *MATa/MATalpha elg1::HIS3MX6/elg1::HIS3MX6*<br>*wpl1::TRP1/wpl1::TRP1 PDS5/pds5::NatMX4 SCC2/SCC2-*<br>*6HA::HPHNT1* | This study |
| HY11121 | *MATa/MATalpha elg1::HIS3MX6/elg1::HIS3MX6*<br>*wpl1::TRP1/wpl1::TRP1 PDS5/pds5::NatMX4 SCC2/scc2-pip-*<br>*6HA::HPHNT1* | This study |
| HY11119 | *MATa/MATalpha elg1::HIS3MX6/elg1::HIS3MX6*<br>*wpl1::TRP1/wpl1::TRP1 PDS5/pds5::NatMX4 SCC2/scc2-20-*<br>*6HA::HPHNT1* | This study |
| HY9940 | *MATa elg1::HIS3MX6 wpl1::TRP1* | This study |
| HY11212 | *MATa elg1::HIS3MX6 wpl1::TRP1 SCC2-6HA::HPHNT1* | This study |
| HY11213 | *MATa elg1::HIS3MX6 wpl1::TRP1 scc2-pip-6HA::HPHNT1* | This study |
| HY11214 | *MATa elg1::HIS3MX6 wpl1::TRP1 scc2-20-6HA::HPHNT1* | This study |
| HY11218 | *MATalpha elg1::HIS3MX6 wpl1::TRP1 SCC2-6HA::HPHNT1* | This study |
| HY11219 | *MATalpha elg1::HIS3MX6 wpl1::TRP1 scc2-pip-6HA::HPHNT1* | This study |
| HY11220 | *MATalpha elg1::HIS3MX6 wpl1::TRP1 scc2-20-6HA::HPHNT1* | This study |
| HY11289 | *MATa/MATalpha elg1::HIS3MX6/elg1::HIS3MX6*<br>*wpl1::TRP1/wpl1::TRP1 CHL1/chl1::KANMX4 SCC2/SCC2-*<br>*6HA::HPHNT1* | This study |
| HY11451 | *MATa/MATalpha CHL1/chl1::KANMX4 SCC2/SCC2*<br>*(tSCC2::HPHNT1)* | This study |
| HY11452 | *MATa/MATalpha CHL1/chl1::KANMX4 SCC2/scc2-pip::HPHNT1* | This study |
| HY11454 | *MATa/MATalpha CTF18/ctf18::KANMX4 SCC2/scc2-pip::HPHNT1* | This study |
| HY11456 | *MATa/MATalpha CTF19/ctf19::KITRP1 SCC2/scc2-pip::HPHNT1* | This study |
| HY11459 | *MATa/MATalpha elg1::HIS3MX6/elg1::HIS3MX6 wpl1::TRP1/wpl1::TRP1 PDS5/pds5::NatMX4 SCC2/SCC2 (tSCC2::HPHNT1)* | This study |
| HY11460 | *MATa/MATalpha elg1::HIS3MX6/elg1::HIS3MX6 wpl1::TRP1/wpl1::TRP1 PDS5/pds5::NatMX4 SCC2/scc2-pip::HPHNT1* | This study |
| HY11575 | *MATa/MATalpha MRC1/mrc1::KANMX4 SCC2/scc2-pip::HPHNT1* | This study |
| HY11602 | *MATa/MATalpha CTF4/ctf4::TRP1 SCC2/scc2-pip::HPHNT1* | This study |
| HY11603 | *MATa/MATalpha CSM3/csm3::HIS3MX6 SCC2/scc2-pip::HPHNT1* | This study |
| HY11604 | *MATa/MATalpha TOF1/tof1::KANMX6 SCC2/scc2-pip::HPHNT1* | This study |
| HY9980 | *MATalpha elg1::HIS3MX6 wpl1::TRP1 pds5::NatMX4* | This study |
| HY11379 | *MATa elg1::HIS3MX6 wpl1::TRP1 SCC2 (tSCC2::HPHNT1)* | This study |
| HY11382 | *MATa elg1::HIS3MX6 wpl1::TRP1 scc2-pip::HPHNT1* | This study |
| HY11461 | *MATa elg1::HIS3MX6 wpl1::TRP1 SCC2 (tSCC2::HPHNT1) pds5::NatMX4* | This study |
| HY11463 | *MATa elg1::HIS3MX6 wpl1::TRP1 scc2-pip::HPHNT1 pds5::NatMX4* | This study |
| HY11377 | *MATa SCC2 (tSCC2::HPHNT1)* | This study |
| HY11380 | *MATa scc2-pip::HPHNT1* | This study |
| HY2192 | *MATalpha chl1::KANMX4* | Lab collection |
| HY11309 | *MATalpha ctf18::KANMX4* | This study |
| HY11301 | *MATalpha ctf19::KITRP1* | This study |
| HY11427 | *MATa SCC2 (tSCC2::HPHNT1) chl1::KANMX4* | This study |
| HY11429 | *MATa SCC2 (tSCC2::HPHNT1) ctf18::KANMX4* | This study |
| HY11435 | *MATa scc2-pip::HPHNT1 chl1::KANMX4* | This study |
| HY11437 | *MATa scc2-pip::HPHNT1 ctf18::KANMX4* | This study |
| HY11630 | *MATa scc2-pip::HPHNT1 ctf4::TRP1* | This study |
| HY11632 | *MATa scc2-pip::HPHNT1 csm3::HIS3MX6* | This study |
| HY11634 | *MATa scc2-pip::HPHNT1 tof1::KANMX6* | This study |
| HY11580 | *MATa scc2-pip::HPHNT1 mrc1::KANMX4* | This study |
| HY6823 | *MATalpha mrc1::KANMX4* | This study |
| HY1613 | *MATalpha ctf4::TRP1* | Lab collection |
| HY2844 | *MATalpha tof1::KANMX6* | Lab collection |
| HY4479 | *MATa csm3::HIS3MX6* | Lab collection |
| HY11479 | *MATa his3-11,15::HIS3tetR-GFP (single integrant), ura3::3xURA3tetO112 SCC2 (tSCC2::HPHNT1)* | This study |
| HY11480 | *MATa his3-11,15::HIS3tetR-GFP (single integrant), ura3::3xURA3tetO112 SCC2 (tSCC2::HPHNT1) chl1::TRP1* | This study |
| HY11481 | *MATa his3-11,15::HIS3tetR-GFP (single integrant), ura3::3xURA3tetO112 scc2-pip::HPHNT1* | This study |
| HY11483 | *MATa his3-11,15::HIS3tetR-GFP (single integrant), ura3::3xURA3tetO112 scc2-pip::HPHNT1 chl1::TRP1* | This study |
| HY11548 | *MATa his3-11,15::HIS3tetR-GFP (single integrant), ura3::3xURA3tetO112 SCC2 (tSCC2::HPHNT1) ctf18::TRP1* | This study |
| HY11549 | *MATa his3-11,15::HIS3tetR-GFP (single integrant), ura3::3xURA3tetO112 scc2-pip::HPHNT1 ctf18::NatNT2* | This study |
| HY11489 | *MATa/MATalpha SCC2 (tSCC2::HPHNT1)/SCC2 (tSCC2::HPHNT1) CTF18/ctf18::KANMX4 CTF19/ctf19::KITRP1* | This study |
| HY11490 | *MATa/MATalpha scc2-pip::HPHNT1/scc2-pip::HPHNT1* | This study |
|  | *CTF18/ctf18::KANMX4 CTF19/ctf19::KITRPI* |  |
| HY11431 | *MATa SCC2 (tSCC2::HPHNT1) ctf19::KITRPI* | This study |
| HY11439 | *MATa scc2-pip::HPHNT1 ctf19::KITRPI* | This study |
| HY11509 | *MATa SCC2 (tSCC2::HPHNT1) ctf18::KANMX4 ctf19::KITRPI* | This study |
| HY11511 | *MATa scc2-pip::HPHNT1 ctf18::KANMX4 ctf19::KITRPI* | This study |
| HY12024 | *MATa/MATalpha SCC2/scc2-E822K::HPHNT1*<br>*SCC4/(KANMX4)pGALS-SCC4* | This study |
| HY12025 | *MATa/MATalpha SCC2/scc2-E822K-pip::HPHNT1*<br>*SCC4/(KANMX4)pGALS-SCC4* | This study |
| HY12037 | *MATa/MATalpha SCC2/scc2-E822K-pol32PIP::NatNT2*<br>*SCC4/(KANMX4)pGALS-SCC4* | This study |
| HY12038 | *MATa/MATalpha SCC2/scc2-E822K-pol32pip::NatNT2*<br>*SCC4/(KANMX4)pGALS-SCC4* | This study |
| HY12039 | *MATa/MATalpha SCC2/scc2-E822K-17aa::NatNT2*<br>*SCC4/(KANMX4)pGALS-SCC4* | This study |
| HY12040 | *MATa/MATalpha SCC2/scc2-E822K-pip-pol32PIP::NatNT2*<br>*SCC4/(KANMX4)pGALS-SCC4* | This study |
| HY11750 | *MATalpha scc2-E822K::HPHNT1 (KANMX4)pGALS-SCC4* | This study |
| HY11746 | *MATalpha scc2-E822K-pip::HPHNT1 (KANMX4)pGALS-SCC4* | This study |
| HY11999 | *MATa (natNT2)pADH1-7His8FLAG-scc2-E822K::HPHNT1* | This study |
| HY11971 | *MATa (natNT2)pADH1-7His8FLAG-scc2-E822K-pip::HPHNT1* | This study |
| HY12139 | *MATa pSCC2-7His8FLAG-scc2-E822K::natNT2* | This study |
| HY12134 | *MATa pSCC2-7His8FLAG-scc2-E822K-pip::natNT2* | This study |
| HY12136 | *MATa pSCC2-7His8FLAG-scc2-E822K-pol32PIP::NatNT2* | This study |
| HY12137 | *MATa pSCC2-7His8FLAG-scc2-E822K-pol32pip::NatNT2* | This study |
| HY11937 | *MATalpha (natNT2)pADH1-7His10FLAG-scc2-E822K::HPHNT1*<br>*(KANMX4)pGALS-SCC4* | This study |
| HY12047 | *MATa POL30-3MYC::KANMX4* | This study |
| HY12089 | *MATa/MATalpha POL30/POL30-3MYC::KANMX4*<br>*SCC2/scc2-E822K::HPHNT1 SCC4/scc4::NatNT2* | This study |
| HY12090 | *MATa scc2-E822K::HPHNT1 scc4::NatNT2* | This study |
| HY12093 | *MATalpha scc2-E822K::HPHNT1 scc4::NatNT2* | This study |
| HY12095 | *MATalpha POL30-3MYC::KANMX4 scc2-E822K::HPHNT1*<br>*scc4::NatNT2* | This study |
| HY12144 | *MATa/MATalpha pol30-6/POL30-3MYC::KANMX4*<br>*scc2-E822K::HPHNT1/scc2-E822K::HPHNT1 SCC4/scc4::NatNT2* | This study |
| HY12150 | *MATa/MATalpha pol30-79/POL30-3MYC::KANMX4*<br>*scc2-E822K::HPHNT1/scc2-E822K::HPHNT1 SCC4/scc4::NatNT2* | This study |
| HY12147 | *MATa/MATalpha pol30-8/POL30-3MYC::KANMX4*<br>*scc2-E822K::HPHNT1/scc2-E822K::HPHNT1 SCC4/scc4::NatNT2* | This study |
| HY12151 | *MATa pol30-79 scc2-E822K::HPHNT1 scc4::NatNT2* | This study |
| HY12048 | *MATa (natNT2)pADH1-7His8FLAG-scc2-E822K::HPHNT1*<br>*POL30-3MYC::KANMX4* | This study |
| HY12050 | *MATa (natNT2)pADH1-7His8FLAG-scc2-E822K-pip::HPHNT1*<br>*POL30-3MYC::KANMX4* | This study |
| HY11980 | *MATa/MATalpha SCC2/scc2-E822K::HPHNT1*<br>*SCC4/(KANMX4)pGALS-SCC4 CTF18/ctf18::TRP1* | This study |
| HY12061 | *MATa/MATalpha SCC2/scc2-E822K::HPHNT1*<br>*SCC4/(KANMX4)pGALS-SCC4 CHL1/chl1::URA3* | This study |
| HY11979 | *MATa/MATalpha SCC4/(KANMX4)pGALS-SCC4*<br>*CTF18/ctf18::TRP1 SCC2/(natNT2)pADH1-7His8FLAG-scc2-*<br>*E822K::HPHNT1* | This study |
| HY12071 | *MATa/MATalpha SCC4/(KANMX4)pGALS-SCC4 CHL1/chl1::URA3*<br>*SCC2/(natNT2)pADH1-7His8FLAG-scc2-E822K::HPHNT1* | This study |
| HY11964 | *MATa/MATalpha SCC4/(KANMX4)pGALS-SCC4 SCC2/(natNT2)pADH1-7His8FLAG-scc2-E822K-pip::HPHNT1* | This study |
| HY12026 | *MATa/MATalpha SCC4/(KANMX4)pGALS-SCC4 CTF18/ctf18::TRP1 SCC2/(natNT2)pADH1-7His8FLAG-scc2-E822K-pip::HPHNT1* | This study |
| HY12062 | *MATa/MATalpha SCC2/scc2-E822K::HPHNT1 SCC4/(KANMX4)pGALS-SCC4 CHL1/chl1-K48R-3HA::LEU2* | This study |
| HY12072 | *MATa/MATalpha SCC4/(KANMX4)pGALS-SCC4 CHL1/chl1-K48R-3HA::LEU2 SCC2/(natNT2)pADH1-7His8FLAG-scc2-E822K::HPHNT1* | This study |
| HY12027 | *MATa/MATalpha CTF18/ctf18::TRP1 CHL1/chl1::KANMX4 SCC2/(natNT2)pADH1-7His8FLAG-scc2-E822K::HPHNT1* | This study |
| HY12105 | *MATa/MATalpha CTF18/ctf18::TRP1 CHL1/chl1-K48R-3HA::LEU2 (natNT2)pADH1-7His8FLAG-scc2-E822K::HPHNT1/(natNT2)pADH1-7His8FLAG-scc2-E822K::HPHNT1* | This study |
| HY12104 | *MATa/MATalpha CTF18/ctf18::TRP1 CHL1/chl1-K48R-3HA::LEU2 (natNT2)pADH1-7His8FLAG-scc2-E822K::HPHNT1/(natNT2)pADH1-7His8FLAG-scc2-E822K::HPHNT1 (KANMX4)pGALS-SCC4/(KANMX4)pGALS-SCC4* | This study |
| HY12153 | *MATa/MATalpha SCC2/pSCC2-7His8FLAG-scc2-E822K::natNT2 SCC4/(KANMX4)pGALS-SCC4* | This study |
| HY12154 | *MATa/MATalpha SCC2/pSCC2-7His8FLAG-scc2-E822K-pip::natNT2 SCC4/(KANMX4)pGALS-SCC4* | This study |
| HY12155 | *MATa/MATalpha SCC2/pSCC2-7His8FLAG-scc2-E822K-pol32PIP::NatNT2 SCC4/(KANMX4)pGALS-SCC4* | This study |
| HY12156 | *MATa/MATalpha SCC2/pSCC2-7His8FLAG-scc2-E822K-pol32pip::NatNT2 SCC4/(KANMX4)pGALS-SCC4* | This study |
| HY12171 | *MATalpha pSCC2-7His8FLAG-scc2-E822K::natNT2* | This study |
| HY12172 | *MATalpha pSCC2-7His8FLAG-scc2-E822K-pip::natNT2* | This study |
| HY12173 | *MATalpha pSCC2-7His8FLAG-scc2-E822K-pol32PIP::NatNT2* | This study |
| HY12174 | *MATalpha pSCC2-7His8FLAG-scc2-E822K-pol32pip::NatNT2* | This study |
| HY11952 | *MATalpha (natNT2)pADH1-7His8FLAG-scc2-E822K::HPHNT1 (KANMX4)pGALS-SCC4* | This study |
| HY11972 | *MATa (natNT2)pADH1-7His8FLAG-scc2-E822K-pip::HPHNT1 (KANMX4)pGALS-SCC4* | This study |
| HY12450 | *MATa/MATalpha SCC1/(TRP1)pGAL1-SCC1 ELG1/elg1::HPHMX6 WPL1/wpl1::HIS3MX6 PDS5/pds5::NatMX4* | This study |
| HY12492 | *MATa (TRP1)pGAL1-SCC1* | This study |
| HY12493 | *MATa (TRP1)pGAL1-SCC1 elg1::HPHMX6* | This study |
| HY12494 | *MATa (TRP1)pGAL1-SCC1 elg1::HPHMX6 pds5::NatMX4* | This study |

All strains are isogenic to W303 background.

**Supplementary Table 2.**
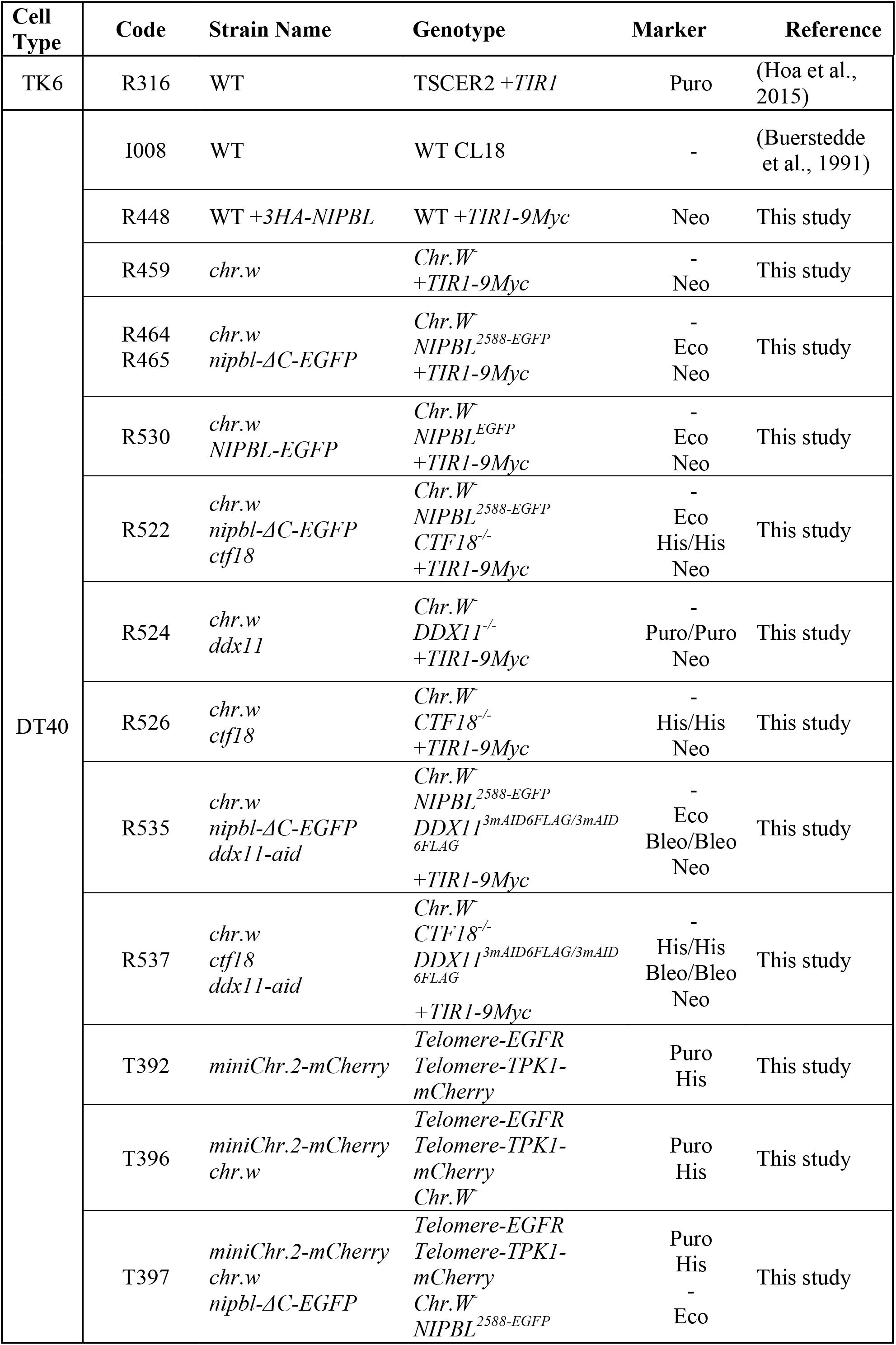
Cell lines used in this study.

## Extended Data Figure Legends

**Extended Data Figure 1:**
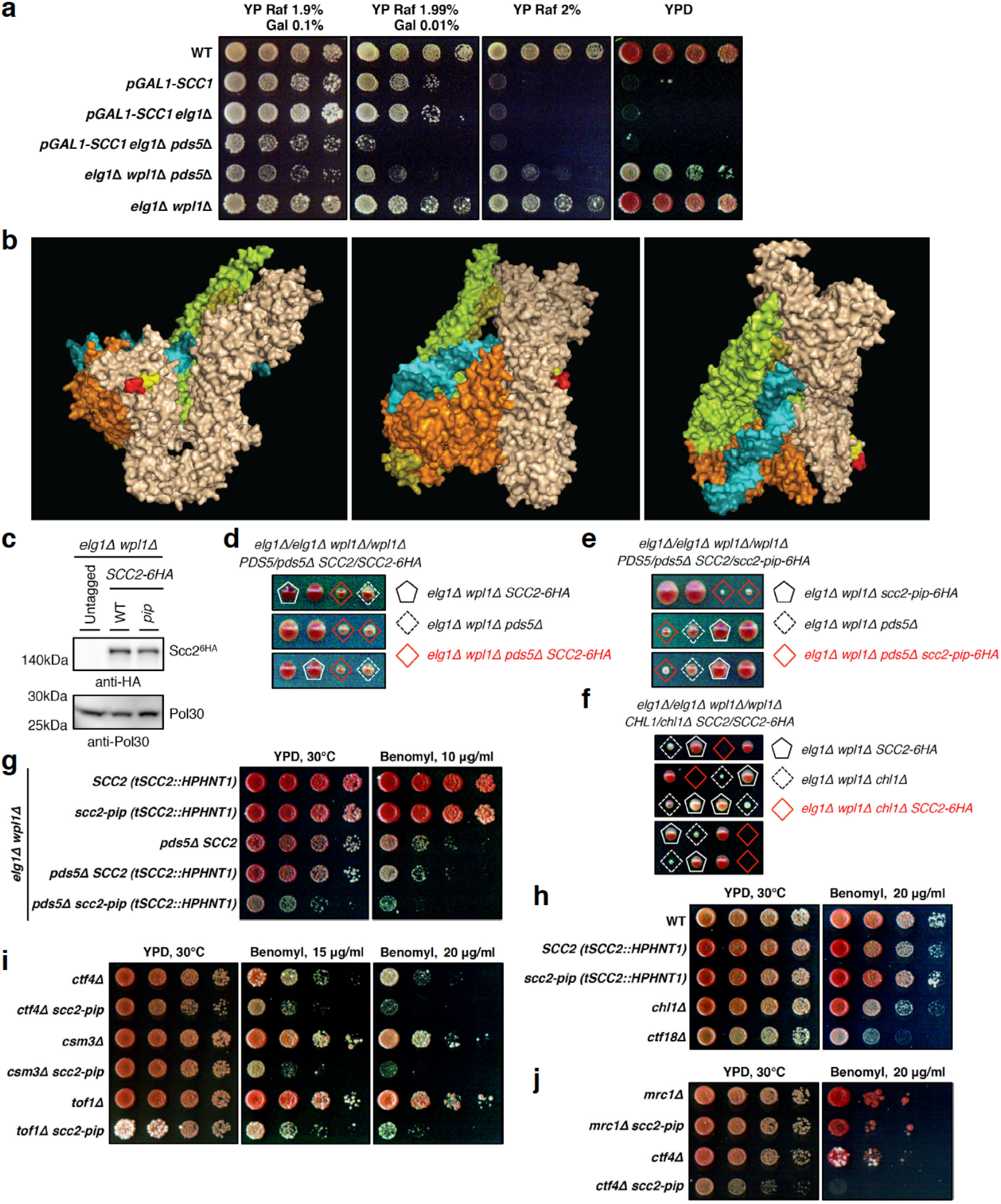
The *scc2-pip* mutant shows additive cohesion defects with replisome mutants of the cohesin conversion, but not de novo loading pathway. **a**, Overexpression of *SCC1* from galactose-inducible *pGAL1* promoter (Gal 0.1%) supports viability of *elg1Δ pds5Δ* cells, whereas low *SCC1* expression (Gal 0.01%) that supports viability of *elg1Δ* cells does not permit the growth of *elg1Δ pds5Δ*mutant. Spotting of 1:7 serial dilutions on indicate plates containing glucose (YPD), raffinose (Raf) and galactose (Gal). **b**, Cryo-EM structure of the budding yeast cohesin-Scc2-DNA complex (PDB 6ZZ6) with C-terminal residues 1466-1470 and 1471-1475 of Scc2 highlighted yellow and red, respectively. The last 18 residues of Scc2 containing consensus PIP motif are not resolved in the structure. The DNA is colored blue, Smc1 – orange, Smc3 – green, Scc1 – olive. **c**, Protein levels of C-terminally 6HA-tagged Scc2 and its PIP mutant variant in *elg1Δ pds5Δ* cells. PCNA (yeast Pol30) served as loading control. **d**-**f**, Tetrad dissection analysis of the indicated diploid strains. C-terminal 6HA-tagging of Scc2 results in the slow-growth phenotype in *elg1Δ wpl1Δ pds5Δ* (**d**), and lethality in *elg1Δ wpl1Δ chl1Δ* (**f**) background. **g**, *scc2-pip* mutant combined with *elg1Δ wpl1Δ pds5Δ* results in slow-growth phenotype and strong sensitivity to benomyl. Strains with disruption of the *SCC2* terminator by selection marker (*tSCC2::HPHNT1*) without mutating the *SCC2* PIP served as controls. **h**-**j**, The *scc2-pip* mutant, while on its own not being sensitive to microtubule poison (**h**), shows strong sensitivity to benomyl when combined with replisome mutants of the cohesin conversion pathway *ctf4Δ, csm3Δ*, and *tof1Δ* (**i**), but not with the de novo cohesin loading pathway mutant *mrc1Δ* (**j**).

**Extended Data Figure 2:**
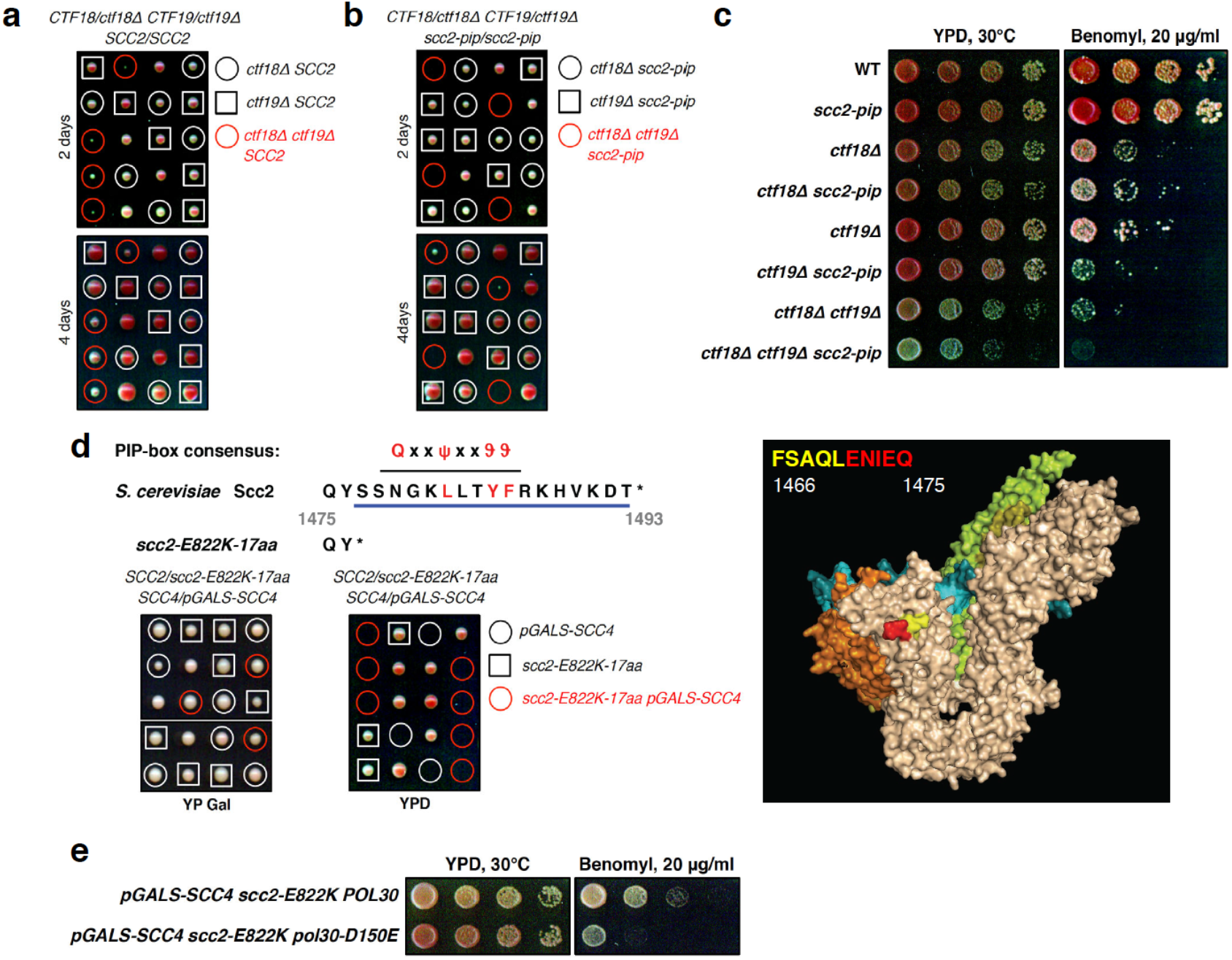
The *scc2-pip* mutant shows additive cohesion defects with *ctf19Δ*, the mutant of kinetochore receptor for cohesin loader, and is synthetic lethal with *scc4Δ*. **a**-**b**, Tetrad dissection analysis of the indicated diploid strains. The *scc2-pip* mutant shows synthetic growth defect when combined with *ctf18Δ ctf19Δ* (**b**) in comparison with *ctf18Δ ctf19Δ* double mutant (**a**). **c**, *scc2-pip* mutant similarly to *ctf18Δ* shows additive sensitivity to benomyl when combined with *ctf19Δ*. **d**, Tetrad dissection analysis of the *SCC2*/*scc2-E822K-17aa SCC4*/*pGALS-SCC4* diploid strain on galactose- (YP Gal) and glucose-containing (YPD) plates. *scc2-E822K-17aa* mutant lacking last 17 residues that harbor consensus PIP motif cannot rescue the lethality of cells expressing *SCC4* from the galactose-inducible *pGALS* promoter when tetrads are dissected on YPD plates. **e**, The disassembly-prone PCNA mutant *pol30-D150E* shows additive sensitivity to benomyl when combined with *scc4 scc2-E822K* double mutant.

**Extended Data Figure 3:**
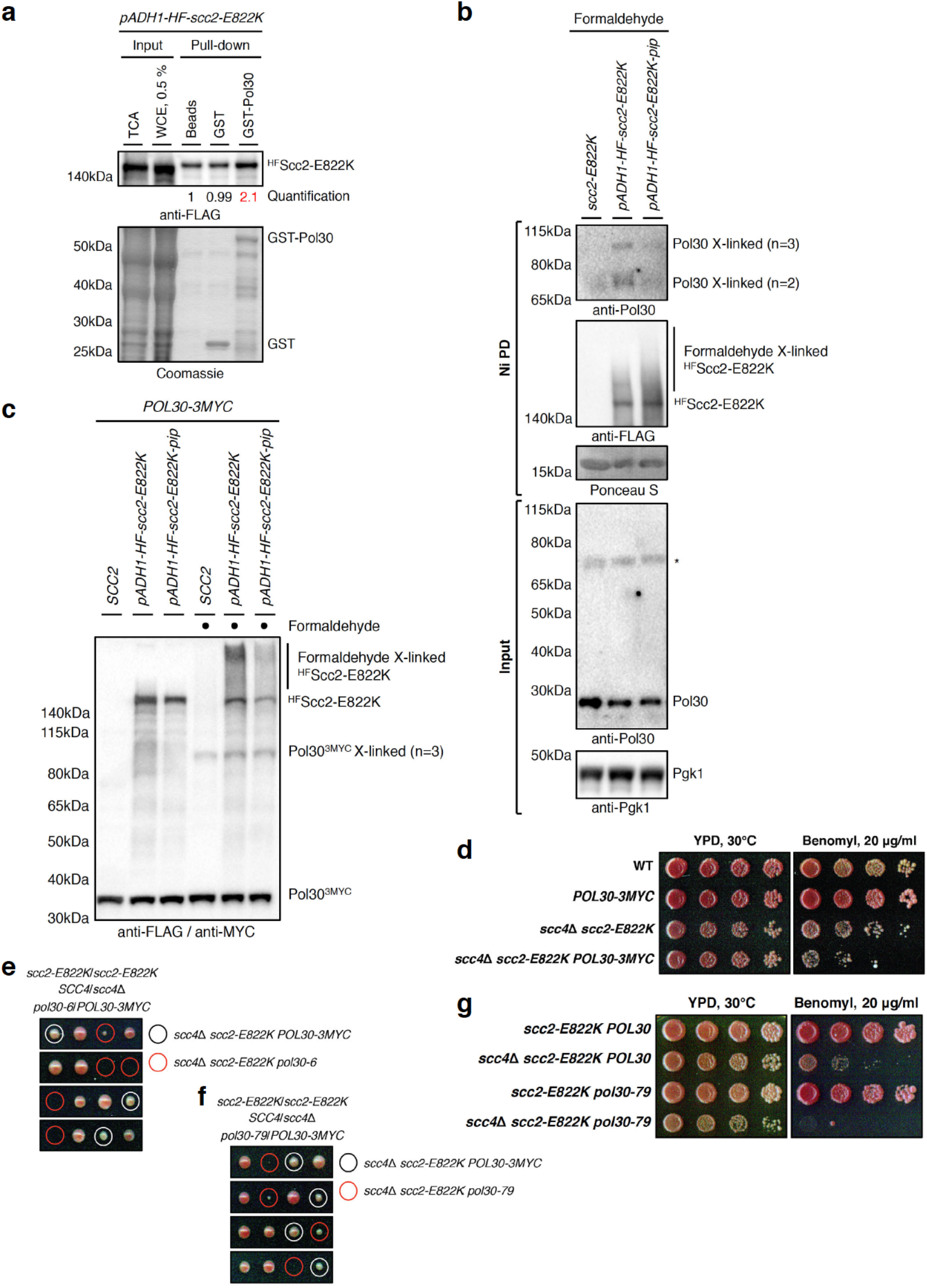
Scc2 interacts with PCNA and *scc4Δ scc2-E822K* double mutant shows additive defects with PCNA mutants. **a**, GST pull-down using recombinant GST-Pol30 (yeast PCNA) in lysates from yeast cells expressing N-terminally 7His8FLAG (HF)-tagged Scc2-E822K from a strong constitutive promoter *pADH1*. **b**, Ni-NTA pull-down (Ni PD) of HF-tagged Scc2-E822K or its PIP mutant under denaturing conditions from yeast cells after formaldehyde crosslinking. ^HF^Scc2-E822K but not its PIP mutant variant binds to cross-linked PCNA species (Pol30 X-linked). PCNA was detected using polyclonal anti-Pol30 antibody (the asterisk indicates a cross-reactive protein), untagged *scc2-E822K* strain was used as control. Ni PD efficiency was assayed using Ponceau S staining (nonspecifically pulled-down protein of ≈15 kDa visualized), Pgk1 served as loading control. **c**, Cross-linked PCNA species (Pol30^3MYC^ X-linked) detected using anti-MYC antibody after formaldehyde crosslinking of yeast cells expressing C-terminally 3MYC-tagged Pol30. **d**, C-terminally 3MYC-tagged PCNA (*POL30-3MYC*) variant shows additive sensitivity to benomyl when combined with *scc4Δ scc2-E822K* double mutant. **e**-**f**, Tetrad dissection analysis of the indicated diploid strains. Mutants causing disruptions of a surface cavity on the front face of the PCNA ring, *pol30-6* (**e**) and *pol30-79* (**f**), show synthetic sickness when combined with *scc4Δ scc2-E822K* double mutant. **g**, *pol30-79* mutant shows additive sensitivity to benomyl when combined with *scc4Δ scc2-E822K* double mutant.

**Extended Data Figure 4:**
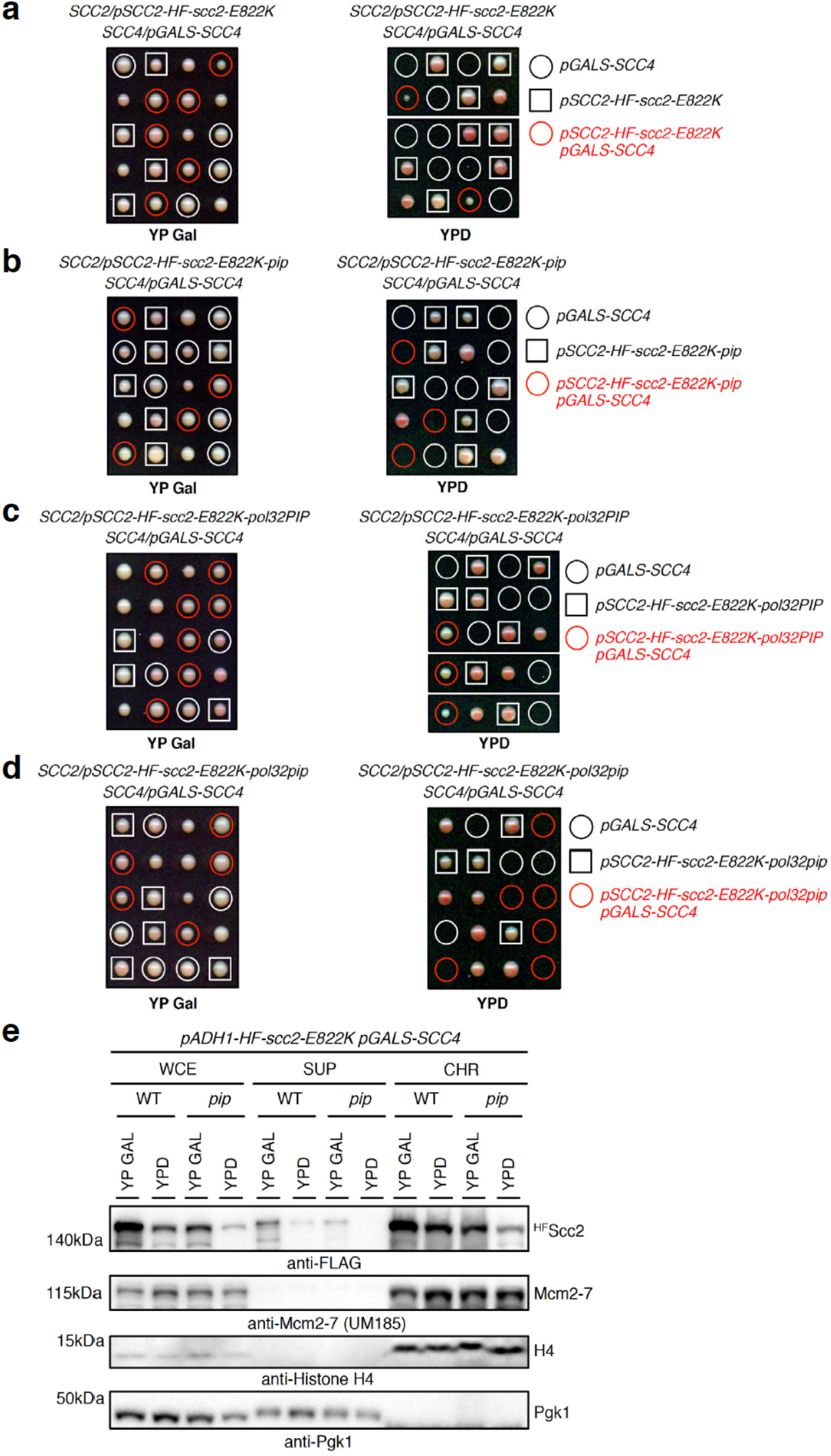
Scc4 and PIP of Scc2 are both contributing to chromatin binding of the cohesin loader. **a**-**d**, Tetrad dissection analysis on the indicated diploid strains on galactose- (YP Gal) and glucose-containing (YPD) plates. N-terminally 7His8FLAG (HF)-tagged Scc2-E822K and its various PIP mutants are expressed from endogenous promoter *pSCC2*. Cohesin loaders ^HF^Scc2-E822K with intact PIP motifs, either endogenous (**a**) or replaced by the PIP of polymerase δ nonessential subunit Pol32 (**c**), support viability upon *SCC4* shut-off on YPD plates. PIP mutants with conserved residues replaced by alanines (**b, d**) on the contrary fail in providing viability. **e**, Subcellular fractionation of cycling cells expressing HF-tagged Scc2-E822K or its PIP mutant from strong constitutive promoter *pADH1* into soluble supernatant (SUP) and chromatin-enriched (CHR) fractions by centrifugation of the whole cell extract (WCE). Chromatin binding of ^HF^Scc2-E822K-pip is decreased compared to ^HF^Scc2-E822K. Used cells in addition had *SCC4* expressed from galactose-inducible *pGALS* promoter allowing to study ^HF^Scc2-E822K chromatin binding upon *SCC4* shut-off after shift from galactose-containing media (YP GAL) to glucose (YPD). Loss of Scc4 further decreases cohesin loader binding to chromatin. To control chromatin fractionation efficiency, the levels of histone H4, replicative helicase Mcm2-7 and the cytoplasmic/plasma membrane protein Pgk1 were detected in fractions.

**Extended Data Figure 5:**
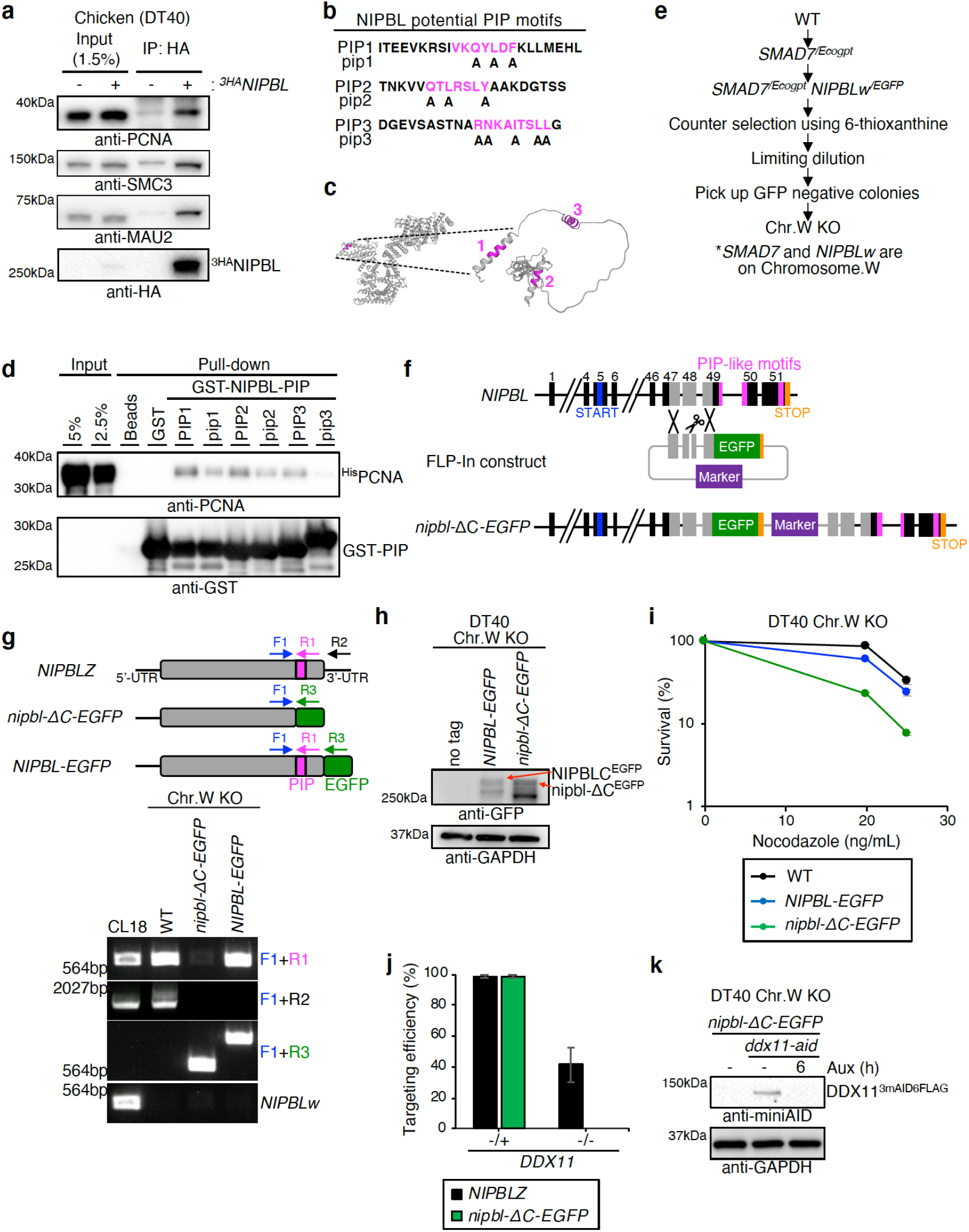
Interaction between PCNA and NIPBL through its PIP-like motifs and establishment of *nipbl-ΔC-EGFP ddx11-aid* conditional mutant. **a**, Immunoprecipitation of overexpressed 3HA-NIPBL in DT40 cells. PCNA was co-immunoprecipitated with 3HA-NIPBL. SMC3 and MAU2 are shown as positive control. **b**, Peptides used for GST in vitro pull-down assay are shown. PIP-like motifs (PIP1-3) at NIPBL C-terminus are colored pink. Amino acids substituted to alanine in the mutant versions of PIPs (pip1-3) are indicated. **c**, Structures of human NIPBL solved by cryo-EM (PDB: 6WGE) and last 203 residues of chicken NIPBL (2581-2783) predicted by AlphaFold2. PIP-like motifs are colored pink. **d**, All three PIP-like motifs fused to GST interact with chicken PCNA. Mutating consensus amino acids of PIP motifs weakens the interaction in vitro. **e**, Scheme of chromosome W knockout establishment in DT40 cells. **f**, Schematic representation of *NIPBL* gene locus on chromosome Z, and FLP-In construct replacing last 195 residues of NIPBL with EGFP. (Closed boxes) exons; (gray boxes) homology arm; (blue box) start codon; (pink boxes) PIP-like motifs; (yellow boxes) stop codon; (green boxes) EGFP; (purple boxes) selection marker gene. **g**, Reverse-transcription PCR analysis verifying *nipbl-ΔC-EGFP* and *NIPBL-EGFP* cells. Primers designed on *NIPBL* cDNA are shown in the upper panel. **h**, Western blotting analysis validating *nipbl-ΔC-EGFP* and *NIPBL-EGFP* cells. **i**, Sensitivity assay using CellTiter-Glo. *NIPBL-EGFP* cells do not exhibit hypersensitivity to nocodazole treatment unlike *nipbl-ΔC-EGFP* cells. **j**, Targeting efficiency of *DDX11* knockout was determined by genomic PCR. At least 18 clones were screened in each experiment. The mean values ± SE are plotted. **k**, Western blotting analysis confirming the depletion of DDX11 tagged with AID, 6 h after auxin addition.

## References

1 Yatskevich, S., Rhodes, J. & Nasmyth, K. Organization of chromosomal DNA by SMC complexes. Annu Rev Genet 53, 445–482 (2019).

2 Uhlmann, F. SMC complexes: from DNA to chromosomes. Nat Rev Mol Cell Biol 17, 399–412 (2016).

3 Onn, I., Heidinger-Pauli, J. M., Guacci, V., Unal, E. & Koshland, D. E. Sister chromatid cohesion: a simple concept with a complex reality. Annu Rev Cell Dev Biol 24, 105–129 (2008).

4 Davidson, I. F. & Peters, J. M. Genome folding through loop extrusion by SMC complexes. Nat Rev Mol Cell Biol 22, 445–464 (2021).

5 Xu, H., Boone, C. & Brown, G. W. Genetic dissection of parallel sister-chromatid cohesion pathways. Genetics 176, 1417–1429 (2007).

6 Srinivasan, M., Fumasoni, M., Petela, N. J., Murray, A. & Nasmyth, K. A. Cohesion is established during DNA replication utilising chromosome associated cohesin rings as well as those loaded de novo onto nascent DNAs. Elife 9, doi:10.7554/eLife.56611 (2020).

7 Petela, N. J. et al. Scc2 is a potent activator of cohesin’s ATPase that promotes loading by binding Scc1 without Pds5. Mol Cell 70, 1134–1148 (2018).

8 Ciosk, R. et al. Cohesin’s binding to chromosomes depends on a separate complex consisting of Scc2 and Scc4 proteins. Mol Cell 5, 243–254 (2000).

9 Kim, Y., Shi, Z., Zhang, H., Finkelstein, I. J. & Yu, H. Human cohesin compacts DNA by loop extrusion. Science 366, 1345–1349 (2019).

10 Murayama, Y. & Uhlmann, F. Biochemical reconstitution of topological DNA binding by the cohesin ring. Nature 505, 367–371 (2014).

11 Collier, J. E. et al. Transport of DNA within cohesin involves clamping on top of engaged heads by Scc2 and entrapment within the ring by Scc3. Elife 9, doi:10.7554/eLife.59560 (2020).

12 Shi, Z., Gao, H., Bai, X. C. & Yu, H. Cryo-EM structure of the human cohesin-NIPBL-DNA complex. Science 368, 1454–1459 (2020).

13 Psakhye, I. & Branzei, D. SMC complexes are guarded by the SUMO protease Ulp2 against SUMO-chain-mediated turnover. Cell Rep 36, 109485 (2021).

14 Murayama, Y. & Uhlmann, F. DNA Entry into and exit out of the cohesin ring by an interlocking gate mechanism. Cell 163, 1628–1640 (2015).

15 Beckouet, F. et al. Releasing activity disengages cohesin’s Smc3/Scc1 interface in a process blocked by acetylation. Mol Cell 61, 563–574 (2016).

16 Liu, H. W. et al. Division of labor between PCNA loaders in DNA replication and sister chromatid cohesion establishment. Mol Cell 78, 725–738 (2020).

17 Zhang, J. et al. Acetylation of Smc3 by Eco1 is required for S phase sister chromatid cohesion in both human and yeast. Mol Cell 31, 143–151 (2008).

18 Sutani, T., Kawaguchi, T., Kanno, R., Itoh, T. & Shirahige, K. Budding yeast Wpl1(Rad61)-Pds5 complex counteracts sister chromatid cohesion-establishing reaction. Curr Biol 19, 492–497 (2009).

19 Ben-Shahar, T. R. et al. Eco1-dependent cohesin acetylation during establishment of sister chromatid cohesion. Science 321, 563–566 (2008).

20 Unal, E. et al. A molecular determinant for the establishment of sister chromatid cohesion. Science 321, 566–569 (2008).

21 Rowland, B. D. et al. Building sister chromatid cohesion: Smc3 acetylation counteracts an antiestablishment activity. Mol Cell 33, 763–774 (2009).

22 Stokes, K., Winczura, A., Song, B., Piccoli, G. & Grabarczyk, D. B. Ctf18-RFC and DNA Pol ε form a stable leading strand polymerase/clamp loader complex required for normal and perturbed DNA replication. Nucleic Acids Res 48, 8128–8145 (2020).

23 Baris, Y., Taylor, M. R. G., Aria, V. & Yeeles, J. T. P. Fast and efficient DNA replication with purified human proteins. Nature, doi:10.1038/s41586-022-04759-1 (2022).

24 Kawasumi, R. et al. Vertebrate CTF18 and DDX11 essential function in cohesion is bypassed by preventing WAPL-mediated cohesin release. Genes Dev 35, 1368–1382 (2021).

25 Moldovan, G. L., Pfander, B. & Jentsch, S. PCNA, the maestro of the replication fork. Cell 129, 665–679 (2007).

26 Michaelis, C., Ciosk, R. & Nasmyth, K. Cohesins: chromosomal proteins that prevent premature separation of sister chromatids. Cell 91, 35–45 (1997).

27 Hinshaw, S. M., Makrantoni, V., Harrison, S. C. & Marston, A. L. The kinetochore receptor for the cohesin loading complex. Cell 171, 72–84 (2017).

28 Munoz, S., Minamino, M., Casas-Delucchi, C. S., Patel, H. & Uhlmann, F. A Role for chromatin remodeling in cohesin loading onto chromosomes. Mol Cell 74, 664–673 (2019).

29 Gerik, K. J., Li, X., Pautz, A. & Burgers, P. M. Characterization of the two small subunits of Saccharomyces cerevisiae DNA polymerase delta. J Biol Chem 273, 19747–19755 (1998).

30 Goellner, E. M. et al. PCNA and Msh2-Msh6 activate an Mlh1-Pms1 endonuclease pathway required for Exo1-independent mismatch repair. Mol Cell 55, 291–304 (2014).

31 Johnson, C., Gali, V. K., Takahashi, T. S. & Kubota, T. PCNA retention on DNA into G2/M phase causes genome instability in cells lacking Elg1. Cell Rep 16, 684–695 (2016).

32 Kondratick, C. M., Litman, J. M., Shaffer, K. V., Washington, M. T. & Dieckman, L. M. Crystal structures of PCNA mutant proteins defective in gene silencing suggest a novel interaction site on the front face of the PCNA ring. PLoS One 13, e0193333 (2018).

33 Kumar, M. et al. ELM-the eukaryotic linear motif resource in 2020. Nucleic Acids Res 48, D296–D306 (2020).

34 Jumper, J. et al. Highly accurate protein structure prediction with AlphaFold. Nature 596, 583–589 (2021).

35 Mirdita, M. et al. ColabFold - making protein folding accessible to all. bioRxiv, doi:10.1101/2021.08.15.456425 (2022).

36 Bhaskara, S. Examination of proteins bound to nascent DNA in mammalian cells using BrdU-ChIP-Slot-Western technique. J Vis Exp, e53647 (2016).

37 Abe, T., Suzuki, Y., Ikeya, T. & Hirota, K. Targeting chromosome trisomy for chromosome editing. Sci Rep 11, 18054 (2021).

38 Nishimura, K., Fukagawa, T., Takisawa, H., Kakimoto, T. & Kanemaki, M. An auxin-based degron system for the rapid depletion of proteins in nonplant cells. Nat Methods 6, 917–922 (2009).

39 Kawasumi, R. et al. ESCO1/2’s roles in chromosome structure and interphase chromatin organization. Genes Dev 31, 2136–2150 (2017).

40 Zheng, G., Kanchwala, M., Xing, C. & Yu, H. MCM2-7-dependent cohesin loading during S phase promotes sister-chromatid cohesion. Elife 7, e33920 (2018).

41 Moldovan, G. L., Pfander, B. & Jentsch, S. PCNA controls establishment of sister chromatid cohesion during S phase. Mol Cell 23, 723–732 (2006).

42 Bender, D. et al. Multivalent interaction of ESCO2 with the replication machinery is required for sister chromatid cohesion in vertebrates. Proc Natl Acad Sci U S A 117, 1081–1089 (2020).

43 Janke, C. et al. A versatile toolbox for PCR-based tagging of yeast genes: new fluorescent proteins, more markers and promoter substitution cassettes. Yeast 21, 947–962 (2004).

44 Sherman, F. Getting started with yeast. Method Enzymol 194, 3–21 (1991).

45 Borges, V. et al. Hos1 deacetylates Smc3 to close the cohesin acetylation cycle. Mol Cell 39, 677–688 (2010).

46 Bowers, J. L., Randell, J. C., Chen, S. & Bell, S. P. ATP hydrolysis by ORC catalyzes reiterative Mcm2-7 assembly at a defined origin of replication. Mol Cell 16, 967–978 (2004).

47 Psakhye, I. & Jentsch, S. Protein group modification and synergy in the SUMO pathway as exemplified in DNA repair. Cell 151, 807–820 (2012).

48 Psakhye, I. & Jentsch, S. Identification of substrates of protein-group SUMOylation. Methods Mol Biol 1475, 219–231 (2016).

49 Psakhye, I., Castellucci, F. & Branzei, D. SUMO-chain-regulated proteasomal degradation timing exemplified in DNA replication initiation. Mol Cell 76, 632–645 (2019).

50 Abe, T. et al. Chromatin determinants of the inner-centromere rely on replication factors with functions that impart cohesion. Oncotarget 7, 67934–67947 (2016).

51 Abe, T. et al. AND-1 fork protection function prevents fork resection and is essential for proliferation. Nat Commun 9, 3091 (2018).

52 Arakawa, H., Lodygin, D. & Buerstedde, J. M. Mutant loxP vectors for selectable marker recycle and conditional knock-outs. BMC Biotechnol 1, 7 (2001).

